# Oxidative phosphorylation safeguards pluripotency via UDP-N-acetylglucosamine

**DOI:** 10.1101/2021.07.06.451381

**Authors:** Jiani Cao, Meng Li, Kun Liu, Xingxing Shi, Ning Sui, Yuchen Yao, Xiaojing Wang, Shaojing Tan, Qian Zhao, Liang Wang, Xiahua Chai, Lin Zhang, Chong Liu, Xing Li, Zhijie Chang, Dong Li, Tongbiao Zhao

## Abstract

The roles of mitochondrial respiration in pluripotency remain largely unknown. We show here that mouse ESC mitochondria possess superior respiration capacity compared to somatic cell mitochondria, and oxidative phosphorylation (OXPHOS) generates the majority of cellular ATP in ESCs. Inhibition of OXPHOS results in extensive pluripotency and metabolic gene expression reprogram, leading to disruption of self-renewal and pluripotency. Metabolomics profiling identifies UDP-N-acetylglucosamine (UDP-GlcNAc) as one of the most significantly decreased metabolites in response to OXPHOS inhibition. The loss of ESC identity induced by OXPHOS inhibition can be ameliorated by directly adding GlcNAc both *in vitro* and *in vivo*. This work demonstrates that mitochondrial respiration, but not glycolysis, produces the majority of ATP in ESCs, and uncovers a novel mechanism whereby mitochondrial respiration is coupled with the hexosamine biosynthesis pathway to generate UDP-GlcNAc for ESC identity maintenance.

**SIGNIFICANCE:** Oxidative phosphorylation (OXPHOS) and glycolysis are the two major pathways for generating ATP in mammalian cells. It is widely assumed that somatic cells utilize OXPHOS, whereas embryonic stem cells (ESCs) utilize glycolysis with low mitochondrial respiration rates even under aerobic conditions. However, the relative contribution of OXPHOS and glycolysis to ATP generation in ESCs, and the role of mitochondrial respiration in regulating ESC identity, have remained unclear. In this study, Cao et al demonstrate that mouse ESC mitochondria have a significantly higher respiration capacity than somatic cell mitochondria. Oxidative phosphorylation produces the majority of cellular ATP in mESCs and is coupled with the hexosamine biosynthesis pathway to generate UDP-GlcNAc for pluripotency maintenance. These findings define the function and mechanism of OXPHOS in regulating pluripotency, and challenge the traditional concept that mESCs rely on glycolysis over OXPHOS for their major supply of energy.

**HIGHLIGHTS:**

1. ESC mitochondria have a significantly higher respiration capability than somatic cell mitochondria
2. OXPHOS, but not glycolysis, produces the majority of cellular ATP in ESCs
3. OXPHOS inhibition induces a decrease in O-GlcNAcylation and the expression of pluripotency genes in blastocysts that can be partially rescued by adding GlcNAc
4. OXPHOS is coupled with the hexosamine biosynthesis pathway for UDP-GlcNAc biosynthesis to maintain ESC identity

## INTRODUCTION

Mitochondria are the metabolic centers of a eukaryotic cell, generating adenosine 5′-triphosphate (ATP) and core metabolites for the biosynthesis of fat, protein and nucleic acids ^1^. In addition, mitochondria function as signaling organelles to communicate with the cytosol to initiate biological events under either homeostatic or stress conditions ^2^. Pluripotent stem cells (PSCs), including embryonic stem cells (ESCs) and induced pluripotent stem cells (iPSCs), have been assumed to possess immature mitochondria and to favor anaerobic glycolysis over oxidative phosphorylation (OXPHOS) for energy production. This proposition is largely based on the findings that ESCs possess globular mitochondria with blurred cristae, and the facts that ESCs have higher glycolysis activity and lower mitochondrial respiration capacity than somatic cells ^3–9^. However, several studies have shown that naïve ESCs have enhanced OXPHOS capacity compared to primed ESCs ^10–12^. In addition, recent studies have shown that mitochondrial autophagy and mitochondrial dynamics are pivotal for ESC self-renewal and pluripotency ^13–17^. These studies have raised a fundamental question: what is the contribution and functional mechanism of mitochondrial respiration in pluripotency regulation?

## RESULTS

### Super-Active Mitochondrial Respiration in mESCs Generates the Majority of Cellular ATP

We first revisited the mitochondrial volume. Due to the imaging limitations of traditional microscopy technology, the precise volume of mitochondria in individual ESCs has not been determined. Taking advantage of advanced resolution structured illumination microscopy (SIM) ^18–20^, we determined the total cellular and mitochondrial volumes of individual mouse naïve ESCs (ESCs), primed ESCs (EpiLCs), neural stem cells (NSCs), embryonic fibroblasts (MEFs) and cardiomyocyte cells (HL-1) (Figures 1A-1C; Videos S1-S5). The total mitochondrial volume in an ESC is similar to that of an EpiLC and significantly smaller than that of an NSC, an MEF and an HL-1 cell (Figure 1B). The cellular volume of an ESC is similar to that of an EpiLC and an HL-1 cell, smaller than an MEF but larger than an NSC (Figure 1C). Consequently, the ratio of mitochondrial volume to whole cell volume in an ESC is significantly smaller than in an EpiLC, an NSC, an MEF or an HL-1 cell (Figure 1D). These new findings prompted us to consider the contribution of mitochondria to ATP generation and stemness regulation in ESCs.

**Figure 1.**
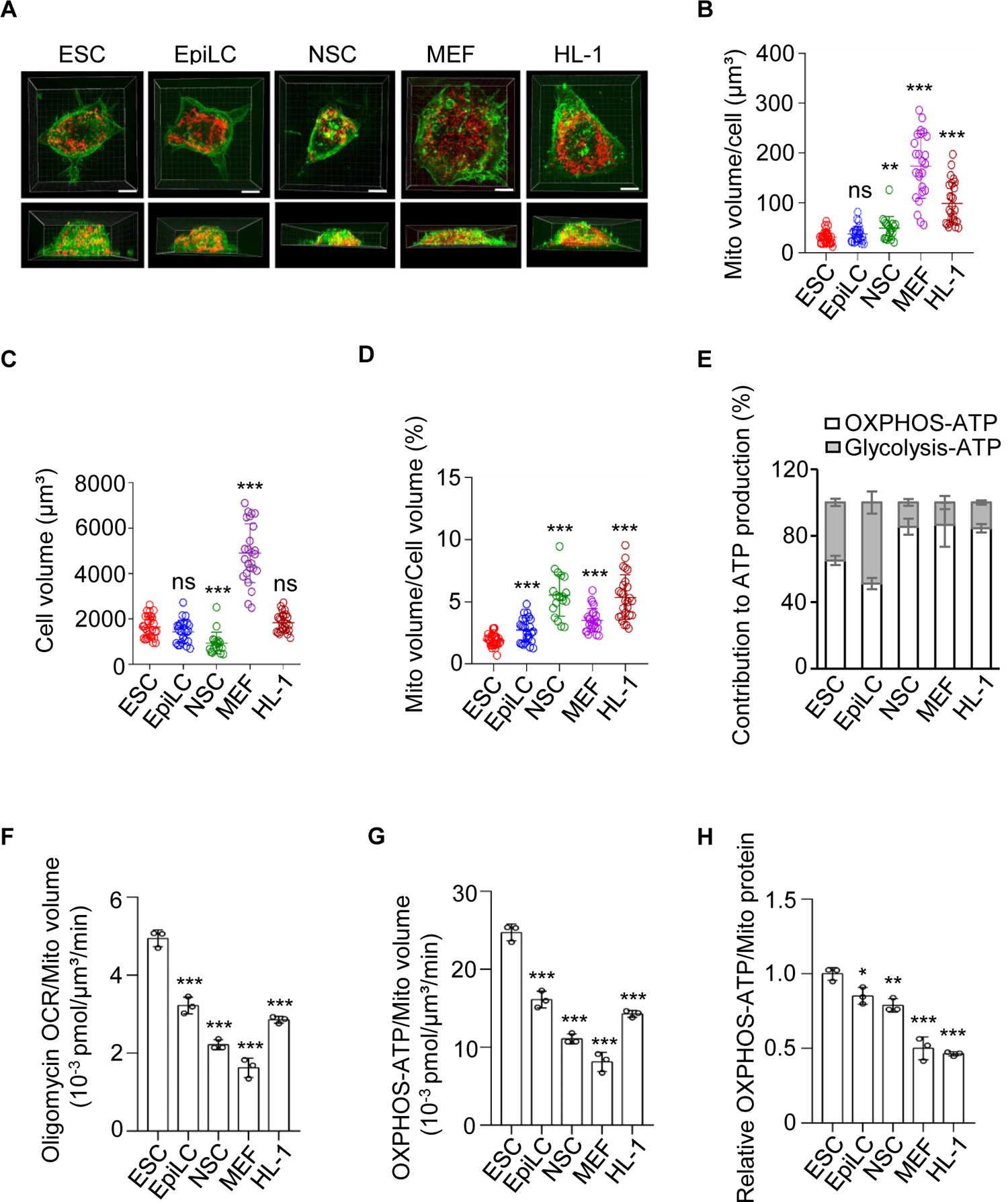
Super-Active Mitochondrial Respiration in mESCs Generates the Majority of Cellular ATP. (**A**) Representative SIM images of a naïve ESC (ESC), a primed ESC (EpiLC), a neural stem cell (NSC), an embryonic fibroblast (MEF) and a cardiomyocyte cell (HL-1). The mitochondria are labeled by mCherry (red) and the cell membranes are stained by DiO (green). Bars, 5 μm. (**B**) The total mitochondrial volume in an ESC (31.05 ± 12.45 µm³) is significantly smaller than that in an NSC (49.02 ± 23.46 µm³), an MEF (174 ± 64.68 µm³) or an HL-1 cell (99.03 ± 42.5 µm³), and is similar to that in an EpiLC (37.4 ± 16.08 µm³). The total mitochondrial volume in individual ESCs, EpiLCs, NSCs, MEFs and HL-1 cells was determined by SIM. Results are shown as mean ± SD; ESC, n=29; EpiLC, n=25; NSC, n=19; MEF, n=25; HL-1, n=26; ^**^ P < 0.01; ^***^P < 0.001; ns, not significant; Student’s *t*-test. (**C**) An ESC (1633 ± 476.7 µm³), which is similar in volume to an EpiLC (1435 ± 491.7 µm³) and an HL-1 cell (1830 ± 416.8 µm³), is smaller than an MEF (4895 ± 1296 µm³) but is larger than an NSC (932.9 ± 476.3 µm³). The volumes of individual ESCs, EpiLCs, NSCs, MEFs and HL-1 cells were determined by SIM. Data are shown as mean ± SD; ESC, n=29; EpiLC, n=25; NSC, n=19; MEF, n=25; HL-1, n=26; ^***^ P < 0.001; ns, not significant; Student’s *t*-test. (**D**) The ratio of total mitochondrial volume to cell volume in ESCs (1.9% ± 0.5%) is smaller than in EpiLCs (2.72% ± 0.98%), NSCs (5.5% ± 1.68%), MEFs (3.5% ± 0.94%) and HL-1 cells (5.36% ± 1.79%). Results are shown as mean ± SD; ESC, n=29; EpiLC, n=25; NSC, n=19; MEF, n=25; HL-1, n=26; ^***^P < 0.001; Student’s *t*-test. (**E**) The relative contribution of OXPHOS and glycolysis to ATP production in ESCs (65.16% ± 2.84% vs 34.84% ± 2.19%), EpiLCs (51.24% ± 3.42% vs 48.76% ± 6.72%), NSCs (85.41% ± 4.86% vs 14.59% ± 2.05%), MEFs (86.62% ± 13.24% vs 13.38% ± 4.00%) and HL-1 cells (84.49% ± 2.59% vs 15.51% ± 1.25%). (**F**) The rate of oligomycin-sensitive oxygen consumption (Oligomycin OCR/Mito volume) is significantly higher in ESCs than in EpiLCs, NSCs, MEFs and HL-1 cells. Values are normalized to mitochondrial volume. Results are shown as mean ± SD of one representative from three independent experiments. n=3; ^***^ P < 0.001; Student’s *t*-test. (**G**) The OXPHOS-ATP generation (OXPHOS-ATP/Mito volume) is significantly higher in ESCs than in EpiLCs, NSCs, MEFs or HL-1 cells. Values are normalized to mitochondrial volume. Results are shown as mean ± SD of one representative from three independent experiments. n=3; ^***^P < 0.001; Student’s *t*-test. (**H**) The OXPHOS-ATP generation (OXPHOS-ATP/Mito mass) is significantly higher in ESCs than in EpiLCs, NSCs, MEFs or HL-1 cells. Values are normalized to mitochondrial protein mass. Results are shown as mean ± SD of one representative from three independent experiments. n=3; ^*^P < 0.05; ^**^P < 0.01; ^***^P < 0.001; Student’s *t*-test.

The oxygen consumption rate (OCR) and extracellular acidification rate (ECAR) were simultaneously measured by a Seahorse XF24 extracellular flux analyzer in ESCs, EpiLCs, NSCs, MEFs and HL-1 cells (Figures S1A and S1B). The absolute quantification of both the oligomycin-sensitive oxygen consumption rate and the glycolytic rate were converted into ATP production rates as previously reported ^21, 22^. Compared to ESCs, EpiLCs and NSCs consumed less oxygen while MEFs and HL-1 cells consumed more oxygen when normalized to equal cell numbers (Figure S1A). Meanwhile, EpiLCs had a higher glycolytic rate than ESCs, and NSCs, MEFs and HL-1 cells had a lower glycolytic rate than ESCs (Figures S1A and S1B). Most strikingly however, OXPHOS generates significantly more ATP than glycolysis in an ESC, an NSC, an MEF and an HL-1 cell. In an EpiLC, OXPHOS and glycolysis generate equal quantities of cellular ATP (Figures 1E and S1C). Similar results were observed in different ESC or iPSC lines (Figures S2A and S2D). When ESCs were cultured in 2i medium, the contribution of OXPHOS to ATP generation further increased (Figures S2B and S2C) ^10, 12^.

Interestingly, when the oxygen consumption rate was normalized to mitochondrial volume, the ESC mitochondria consumed significantly more oxygen than mitochondria in EpiLCs, NSCs, MEFs and even HL-1 cardiomyocytes for ATP-generation-related respiration (Figures 1F and S1K). Correspondingly, ESC mitochondria showed a significantly higher ATP generation capacity than mitochondria in EpiLCs, NSCs, MEFs and HL-1 cells (Figure 1G). To further strengthen this conclusion, mitochondrial mass was used to normalize ATP-generation-related respiration at the same time. The expression of the mitochondrial protein UQCRC2 was used for normalization, as its quantity per microgram of mitochondrial protein was very similar in each tested line, in contrast to other mitochondrial proteins such as TOM40, TIM23 and ATP5A (Figure S1D). For each cell line, the mass of an individual cell was determined and UQCRC2 was used for mitochondrial normalization (Figures S1E-S1H). ESC mitochondria showed the highest ATP generation capacity among all tested cell lines (Figure 1H). In addition, the mitochondrial respiration capacity was determined using cells treated with digitonin and the resultant data were normalized to either mitochondrial volume or mitochondrial mass. Using either normalization parameter, ESC mitochondria showed the biggest ATP generation capacity compared to mitochondria in EpiLCs, NSCs, MEFs and HL-1 cells (Figures S1I-S1L).

These results support the concept that ESC mitochondria stay in an active state with high respiration capacity that generates the majority of cellular ATP. This is opposite to the traditional concept that ESCs rely on glycolysis for their major energy supply ^5, 8, 23^.

### OXPHOS Inhibition Results in More Extensive Gene Expression Reprogramming than Glycolysis Inhibition

To investigate how OXPHOS functions in ESCs, we established assays for dose-dependent inhibition of OXPHOS or glycolysis based on their ATP generation levels. Oligomycin was titrated to inhibit 20% and 50% of the total OXPHOS-ATP generation (designated 20%OI and 50%OI). The oligomycin concentrations for 20%OI and 50%OI were 1 nM and 10 nM, respectively (Figure S3A). Meanwhile, 2-deoxy-D-glucose (2-DG), a glucose analog, was titrated to inhibit 20% and 50% of the total glycolysis-ATP generation (designated 20%GI and 50%GI). The 2-DG concentrations for 20%GI and 50%GI were 5 mM and 10 mM, respectively (Figure S3B). It is worth mentioning that 20%OI and 50%GI have similar inhibition effects on total cellular ATP generation (Figure S3C). Oligomycin at 1 µM and 2-DG at 100 mM completely inhibited OXPHOS-ATP generation and glycolysis-ATP generation respectively (Figures S3A, S3B and S3D).

Transcriptome profiling was employed to investigate gene expression reprogramming in response to OXPHOS or glycolysis inhibition using the titrated concentrations of oligomycin and 2-DG. Surprisingly, OXPHOS inhibition resulted in more significant changes in the gene expression program compared to glycolysis inhibition (Figure 2A; Table S1). In ESCs with 20% inhibition of OXPHOS, there were 4240 differentially expressed genes (DEGs) (1975 down-regulated genes and 2265 up-regulated genes), which is significantly higher than the 1639 DEGs (1007 down-regulated genes and 632 up-regulated genes) in ESCs with 20% inhibition of glycolysis (Figure 2B). The number of DEGs increased to 7510 (3779 down-regulated genes and 3731 up-regulated genes) in ESCs with 50% OXPHOS inhibition. In contrast, 50% inhibition of glycolysis led to 2695 DEGs (1223 down-regulated genes and 1472 up-regulated genes), which is even less than 20% OXPHOS inhibition (Figure 2B). ESCs with 20%OI and 50%OI shared 3919 DEGs, while ESCs with 20%GI and 50%GI shared 1426 DEGs (Figure 2C). These data indicate that OXPHOS inhibition in ESCs results in much more extensive effects on gene expression than glycolysis inhibition at the whole transcriptome level.

**Figure 2.**
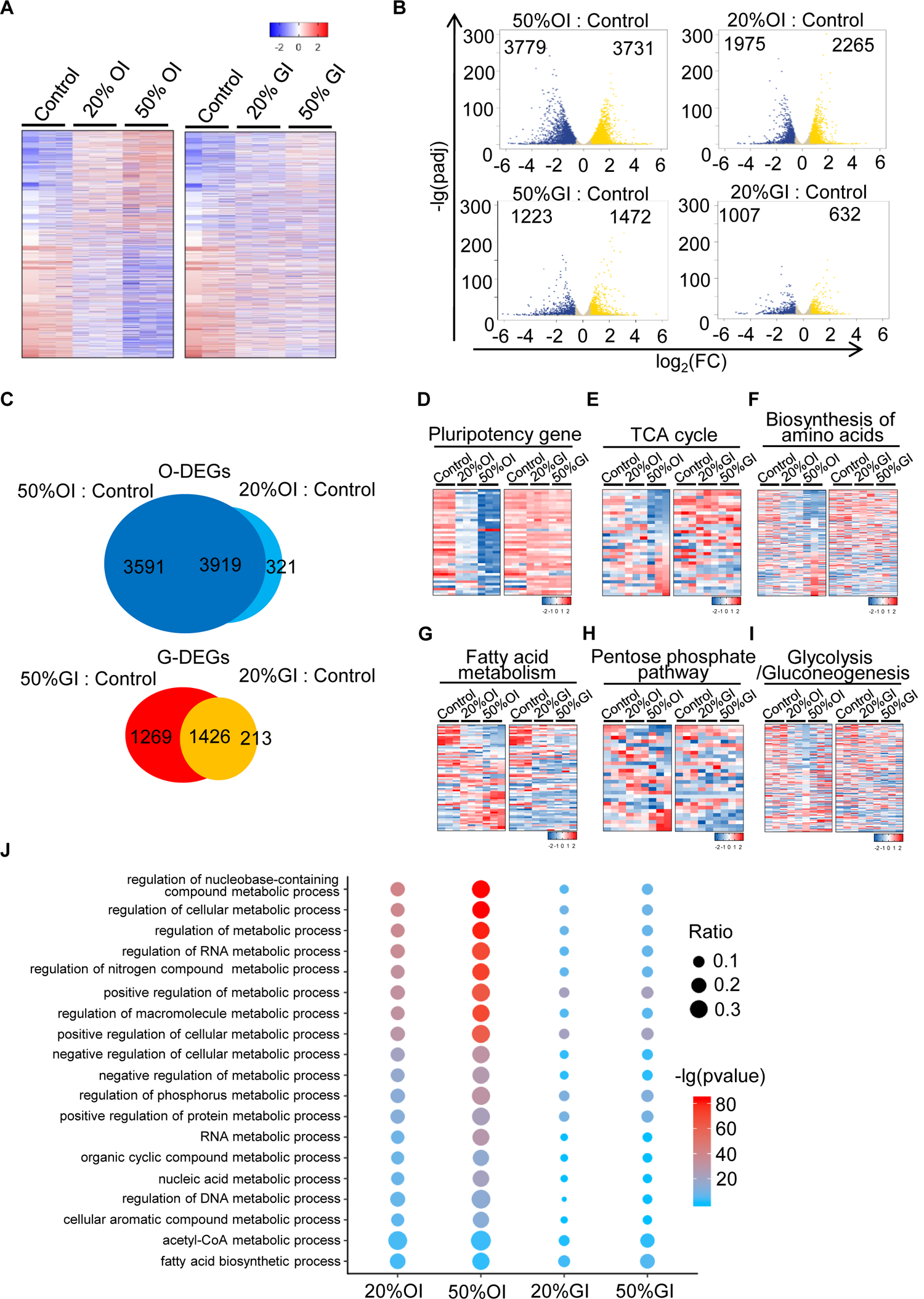
OXPHOS Inhibition Results in More Extensive Gene Expression Reprogramming than Glycolysis Inhibition. (**A**) RNA-seq meta-analysis reveals that OXPHOS inhibition results in dramatic changes of gene expression at the whole-transcriptome level. Values displayed correspond to the expression level in the indicated sample scaled by the mean expression of each gene across samples. (**B**) Volcano plots show the differentially expressed genes (DEGs) between wild-type ESCs and OXPHOS- or glycolysis-inhibited ESCs. (**C**) OXPHOS inhibition results in more DEGs than glycolysis inhibition. The DEGs shared by both 20%OI and 50%OI ESCs are termed as OXPHOS-related differentially expressed genes (O-DEGs), while the DEGs shared by both 20%GI and 50%GI ESCs are termed as glycolysis-related differentially expressed genes (G-DEGs). (**D** to **I**) Inhibition of OXPHOS reprograms expression of pluripotency and metabolic genes. Heatmaps show relative expression levels of genes involved in pluripotency (**D**), TCA pathway (**E**), amino acid biosynthesis (**F**), fatty acid metabolism (**G**), pentose phosphate pathway (**H**), and glycolysis/gluconeogenesis (**I**) upon OXPHOS or glycolysis inhibition. (**J**) Enriched GO term analysis reveals dramatic changes in expression of genes involved in metabolic processes upon OXPHOS inhibition. The gene ratio is indicated by the dot size (the bigger the dot, the greater the ratio) and the significance is indicated by the color of the dot (red, low *P* value; blue, high *P* value). 20%OI, 50%OI: inhibition of ATP production by OXPHOS to 20% or 50% of its maximum; 20%GI, 50%GI: inhibition of ATP production by glycolysis to 20% or 50% of its maximum.

Then we analyzed the expression levels of pluripotency and metabolic genes in these OXPHOS or glycolysis inhibited ESCs. Intriguingly, analysis of a panel of 43 pluripotency genes in oligomycin- or 2-DG-treated ESCs showed that OXPHOS inhibition caused a much stronger downregulatory effect than glycolysis inhibition (Figure 2D). Inhibition of OXPHOS not only disrupted expression of genes in the tricarboxylic acid (TCA) cycle, but also decreased expression of genes in the amino acid biosynthesis, fatty acid metabolism and pentose phosphate pathways as well as the glycolysis/gluconeogenesis pathways in ESCs (Figures 2E-2I). Remarkably, even 20% OXPHOS inhibition has far more extensive effects on the expression of genes in some of these pathways than 50% glycolysis inhibition (Figures 2E-2I). This was further confirmed by GO analysis. The ranked enrichment of GO terms showed more dramatically enhanced clustering of metabolic processes in oligomycin-than 2-DG-treated ESCs (Figure 2J). Thus, mitochondrial respiration serves as a more important pathway than glycolysis for gene expression regulation in ESCs.

### Moderate Inhibition of OXPHOS rather than Glycolysis Disrupts ESC Stemness

Next, the self-renewal and pluripotency of ESCs were tested upon inhibition of mitochondrial respiration or glycolysis. Surprisingly, both 20%OI and 50%OI inhibition significantly decreased ESC colony formation and expression of pluripotency genes, whereas 20%GI did not affect ESC self-renewal and pluripotency, and 50%GI inhibited ESC self-renewal and pluripotency to a lesser extent than 20%OI (Figures 3A-3C and S4A). Importantly, the decreased levels of colony formation resulting from OXPHOS inhibition were partially recovered when oligomycin was withdrawn for different lengths of time (Figure S4B). In addition, treatment with oligomycin or 2-DG at the titrated concentrations for 20%OI, 50%OI, 20%GI and 50%GI did not enhance ESC apoptosis (Figure S3E). These data provide evidence that OXPHOS is essential for ESC self-renewal and pluripotency.

**Figure 3.**
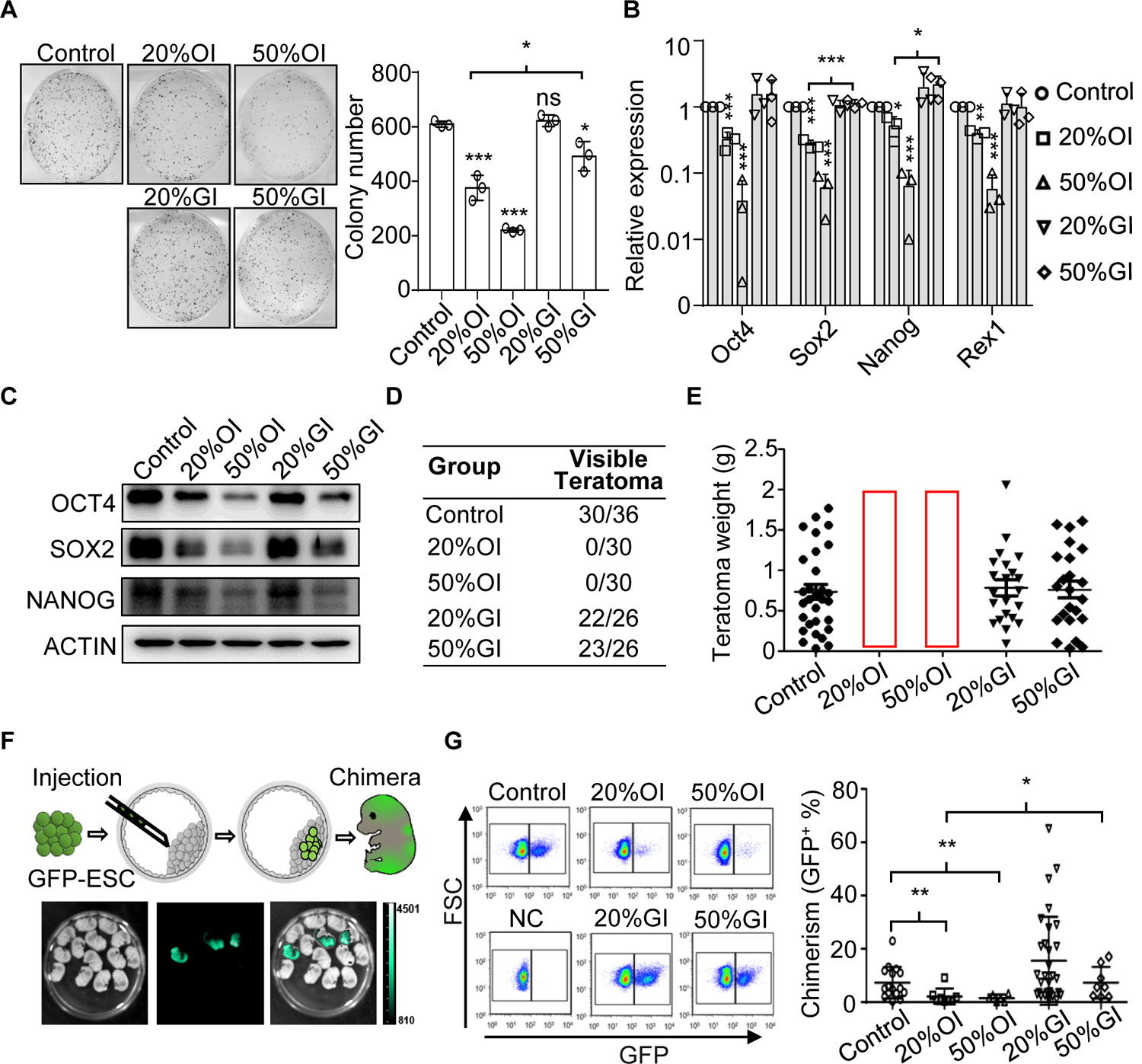
Moderate Inhibition of OXPHOS rather than Glycolysis Disrupts ESC Stemness. (**A**) Moderate inhibition of OXPHOS, but not glycolysis, diminishes the capacity of ESCs for self-renewal. Left, representative images of alkaline phosphatase staining of colonies formed by ESCs treated with either oligomycin or 2-DG. Right, statistical analysis of the number of alkaline phosphatase-positive colonies. Results are shown as mean ± SD from 3 independent experiments. ^*^ P < 0.05; ^***^ P < 0.001; ns, not significant; Student’s *t*-test. (**B**) Inhibition of OXPHOS but not glycolysis decreases mRNA expression of pluripotency genes. Results are shown as mean ± SD from 3 independent experiments. ^*^P < 0.05; ^**^P < 0.01; ^***^P < 0.001; Student’s *t*-test. (**C**) Moderate inhibition of OXPHOS leads to reduced OCT4, SOX2 and NANOG protein expression. Images are representative of 3 independent western blotting experiments. (**D**) Summary of teratoma formation by ESCs with the indicated treatments in nude mice. (**E**) Inhibition of OXPHOS abolishes the teratoma formation capability of ESCs. Each dot represents the weight of a teratoma formed by ESCs with the indicated treatments. Control, n=30; 20%OI, n=30; 50%OI, n=30; 20%GI, n=22; 50%GI, n=23. (**F**) Diagram of the chimeric mouse formation assay. Top, after different treatments, GFP-labeled B6 ESCs were injected into CF1 mouse blastocysts, and the blastocysts were transplanted into surrogate mice. Then the chimeric embryos were isolated and digested into single cells at embryonic day 13.5 (E13.5) for FACS analysis. Bottom, representative images of the chimeric embryos isolated from a surrogate mouse at E13.5. (**G**) Inhibition of OXPHOS, but not glycolysis, decreases the contribution of ESCs to chimeras. Left, GFP-positive cells detected by FACS indicate the number of cells in each chimeric embryo that were derived from the transplanted original cells. Right, summary of data from chimeric embryos. Each dot represents the percentage of GFP^+^ cells in an individual chimeric embryo. Control, n=16; 20%OI, n=8; 50%OI, n=6; 20%GI, n=30; 50%GI, n=8; ^**^P < 0.01; Student’s *t*-test.

The effect of mitochondrial respiration inhibition on ESC differentiation potency was evaluated using teratoma formation and blastocyst chimerism assays. After transplantation into immune-deficient mice, ESCs treated with 20%OI or 50%OI did not form any visible teratomas, while 20%GI or 50%GI had no obvious effects on teratoma formation (Figures 3D, 3E, S4C, and S4D). Accordingly, only a small population of OXPHOS-inhibited ESCs contributed to the chimeric mice, and the chimerism rate with 20%OI or 50%OI ESC was significantly lower than with mock-treated ESCs. In contrast, neither 20%GI nor 50%GI decreased the chimerism rate (Figures 3F and 3G). These data support the notion that OXPHOS guards ESC developmental competence.

Doxycycline-inducible genetic knockdown of ATP5a1, a subunit of the ATP synthase complex, was employed to inhibit mitochondrial function to further clarify the OXPHOS functions in ESC stemness regulation. The ATP5a1 expression level was strictly controlled by the concentration of doxycycline (Figure S5A). Consistent with the chemical treatment results, the mitochondrial respiration, self-renewal, pluripotency and differentiation capability of ESCs were inhibited by ATP5a1 knockdown (Figures S5B and S6). These data confirmed the function of OXPHOS in safeguarding ESC identity.

### OXPHOS Couples with UDP-GlcNAc Generation to Safeguard Pluripotency

Mitochondrial respiration not only contributes to bioenergetics and biosynthesis, but also functions as a signal transduction hub by providing intermediate metabolites as signaling molecules. Considering that OXPHOS inhibition reprograms metabolic gene expression, we asked whether the disruption of ESC self-renewal and pluripotency is attributed to defective metabolite-mediated signal transduction. To this end, targeted profiling of metabolites in the TCA cycle, glycolysis pathway, pentose phosphate pathway, amino acid metabolism and purine metabolism was performed to detect the metabolites that changed in response to OXPHOS inhibition. Interestingly, UDP-N-acetylglucosamine (UDP-GlcNAc), an amino sugar produced by the hexosamine biosynthetic pathway (HBP), was identified at the top of the metabolite list with a dramatic reduction upon inhibition of OXPHOS, and the expression levels of enzymes involved in UDP-GlcNAc biosynthesis were dramatically disturbed (Figures 4A and S7A; Table S2).

**Figure 4.**
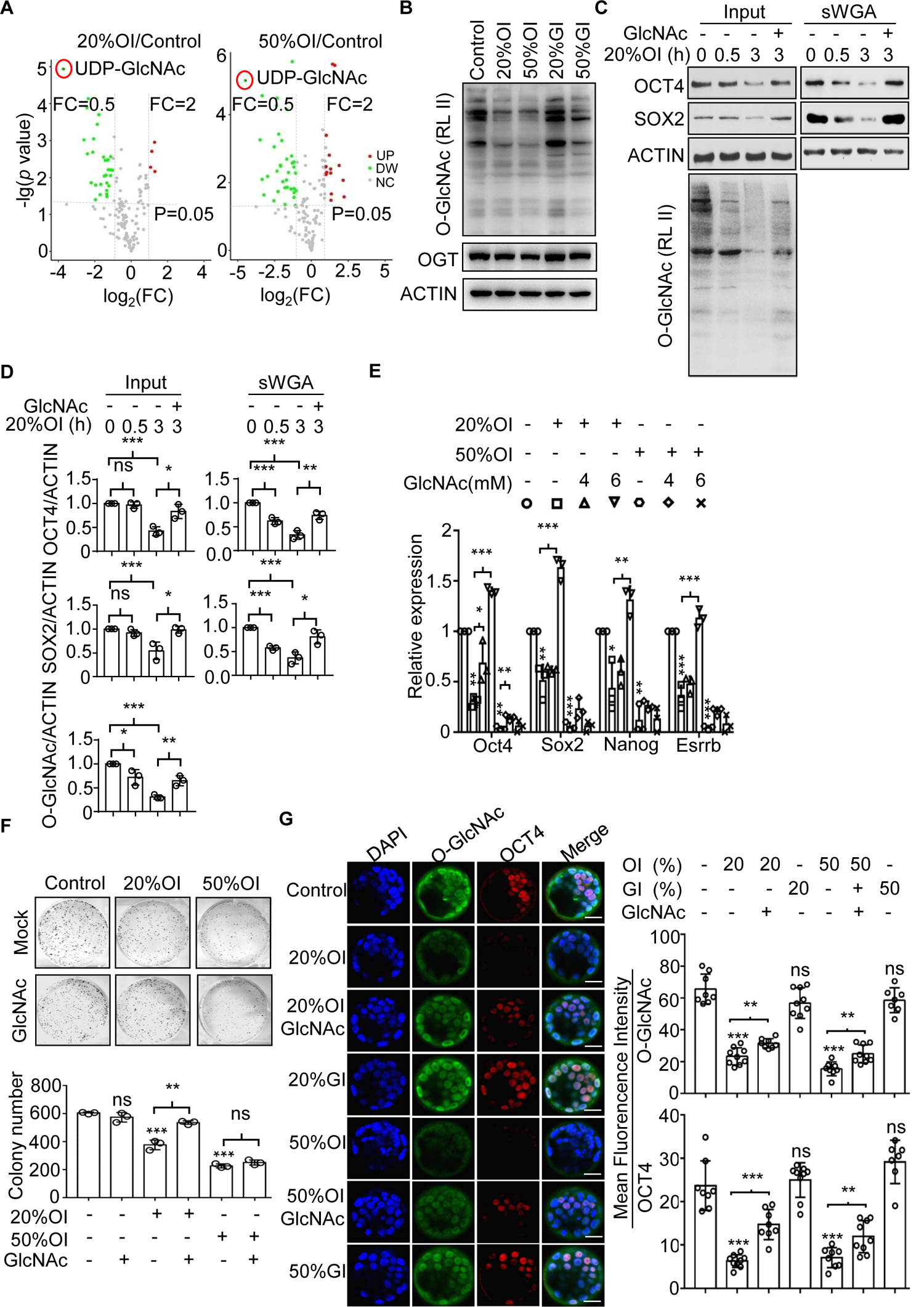
OXPHOS Couples with UDP-GlcNAc Generation to Safeguard Pluripotency. (**A**) Visualization of differential metabolite profiles by Volcano Plots. UDP-GlcNAc (circled in red) is the top down-regulated metabolite upon OXPHOS inhibition. Red dots represent up-regulated metabolites, FC ≥ 2, p ≤ 0.05; green dots represent down-regulated metabolites, FC ≤ 1/2, p ≤ 0.05. p = 0.05 is labeled by the dotted line. FC, Fold change. (**B**) OXPHOS inhibition results in decreased O-GlcNAcylation of cellular proteins independent of OGT expression. O-GlcNAcylation of total cellular protein was detected by western blotting using an anti-O-GlcNAc antibody. Images are representative of 3 independent experiments. (**C**) OXPHOS inhibition causes decreased O-GlcNAcylation and expression of SOX2 and OCT4. These effects are ameliorated by GlcNAc. Decrease of SOX2/OCT4 O-GlcNAcylation initiates 0.5 h after oligomycin addition which is before SOX2/OCT4 downregulation. Cell lysates from ESCs, ESCs treated with oligomycin, or ESCs treated with oligomycin plus GlcNAc for different lengths of time were pulled down by sWGA-agarose and blotted with anti-OCT4, SOX2, O-GlcNAc and β-ACTIN antibodies. (**D**) Statistical analysis of the western blot results shown in **C**. n=3; ^*^P < 0.05; ^**^P < 0.01; ^***^P < 0.001; ns, not significant; Student’s *t*-test. (**E**) OXPHOS inhibition causes decreased transcription of pluripotency genes, which is partially ameliorated by the addition of GlcNAc. Results are shown as mean ± SD from 3 independent experiments. ^*^P < 0.05; ^**^P < 0.01; ^***^P < 0.001; Student’s *t*-test. (**F**) OXPHOS inhibition reduces the colony formation capability of ESCs. This effect is partially ameliorated by the addition of GlcNAc. Top, representative images of ESC colonies stained by alkaline phosphatase; Bottom, statistical analysis of alkaline phosphatase-positive colonies. Results are shown as mean ± SD from 3 independent experiments. ^* *^P < 0.01; ^***^P < 0.001; ns, not significant; Student’s *t*-test. (**G**) Decreased O-GlcNAcylation and expression of OCT4 by OXPHOS inhibition in the inner cell mass of *ex vivo*-cultured blastocysts are recovered by addition of GlcNAc. Left, Immuno-detection was performed with antibodies specific to O-GlcNAc (green) and OCT4 (red), Nuclei were stained with DAPI, Bars, 25 µm; Right, Statistical analysis of the mean fluorescence intensity. Control, n=8; 20%OI, n=9; 20%OI+GlcNAC, n=8; 20%GI, n=9; 50%OI, n=8; 50%OI+GlcNAC, n=9; 50%GI, n=7; ^**^P < 0.01; ^***^P < 0.001; ns, not significant; Student’s t-test.

UDP-GlcNAc serves as the substrate for O-GlcNAcylation of critical regulators of diverse cellular processes, including several known pluripotency factors ^24^. O-GlcNAcylation ensures pluripotency in ESCs by directly regulating the transcriptional activities of key components of the pluripotency network ^25^. The upstream signals that regulate O-GlcNAcylation in ESCs are unknown. Intriguingly, inhibition of OXPHOS led to a dramatic decrease of UDP-GlcNAc and global O-GlcNAcylation, including O-GlcNAcylation of SOX2/OCT4, without affecting O-linked beta-D-N-acetylglucosamine transferase (OGT) expression (Figures 4B, 4C, 4D, S7B, S7C). The decrease of SOX2/OCT4 O-GlcNAcylation began as early as 0.5h after oligomycin addition, prior to downregulation of SOX2/OCT4 expression (Figures 4C, 4D, S7B, S7C). In contrast, glycolysis inhibition had little effect (Figure 4B). In addition, inhibition of UDP-GlcNAc biosynthesis with 6-diazo-5-oxo-L-norleucine (Don) inhibited colony formation and expression of pluripotency genes in ESCs (Figure S7D). Thus, UDP-GlcNAc links OXPHOS to pluripotency.

In addition to *de novo* synthesis, the cellular UDP-GlcNAc pool can be complemented with GlcNAc ^26^ or glucosamine ^27^. To further clarify that OXPHOS regulates ESC identity through UDP-GlcNAc, we added GlcNAc or Glucosamine to oligomycin-treated ESCs to rescue the deterioration in ESC identity. As expected, global protein O-GlcNAcylation as well as the expression of OCT4 and SOX2 were partially restored by adding GlcNAc or glucosamine into OXPHOS-inhibited ESCs (Figures 4C, 4D, S7B, S7C, S7E). Correspondingly, the reduced expression of pluripotency genes and the decreased colony formation capacity in OXPHOS-inhibited ESCs were also compensated by adding GlcNAc (Figures 4E, 4F, S7F). Adding GlcNAc into OXPHOS-inhibited ESCs partially restored the gene expression pattern at the whole transcriptome level as well (Figure S7G). Together, these data support the idea that OXPHOS maintains pluripotency through UDP-GlcNAc.

Importantly, inhibition of OXPHOS, but not glycolysis, resulted in decreased O-GlcNAcylation and expression of Oct4 and Sox2 in the inner cell mass (ICM, the *in vivo* equivalent of ESCs) of blastocysts (Figures 4G, S7H, S7I). The deterioration in pluripotency gene expression in the ICM upon OXPHOS inhibition can be partially rescued by directly supplementing GlcNAc. Thus OXPHOS safeguards pluripotency via UDP-GlcNAc *in vivo*.

## DISCUSSION

Naïve ESCs functionally resemble the inner cell mass of a pre-implantation blastocyst, while primed ESCs resemble post-implantation epiblast cells ^28^. Both mouse and human PSCs possess fewer mitochondria than somatic fibroblast cells and PSC mitochondria have a globular shape and poorly developed cristae. Based on these observations, it was proposed that PSC mitochondria are in an immature and inactive state with low OXPHOS capacity ^3, 29, 30^. However, there is evidence that mitochondria in hPSCs, which are considered to be primed pluripotent cells, possess functional respiration complexes and already consume O_2_ at their maximal capacity ^31^. Other studies have shown that naïve ESC mitochondria can somehow further enhance their OXPHOS capacity compared to mitochondria in primed ESCs ^10–12^. It remains controversial whether the mitochondria in PSCs are in an active or inert state. One of the reasons for this is that the existing studies compared naïve and primed PSCs, or primed PSCs and somatic cells, but they did not directly compare naïve PSCs, primed PSCs and somatic cells. In addition, as PSCs undergo differentiation, many cellular parameters change, like cell volume, cell mass, mitochondrial volume, mitochondrial mass, mtDNA copy number, activity of glyceraldehyde-3-phosphate dehydrogenase (GAPDH), and expression of the mitochondrial house-keeping genes TOM40, TIM23, ATP5A, etc. (Figures 1B, 1C, S1D, S1F) ^3, 31–36^. These changes make it difficult to objectively compare the mitochondrial state in cells at distinct developmental stages. Taking advantage of advanced resolution structured illumination microscopy technology, we determined the absolute volume of individual cells and their mitochondria. The volume of mitochondria in a naïve ESC is significantly lower than that in an NSC, an MEF or an HL-1 cardiomyocyte, and is similar to that in an EpiLC. In terms of cell volume, an ESC is similar to an EPiLC or an HL-1 cell, significantly smaller than an MEF and significantly larger than an NSC (Figure 1A-1C and Movies 1-5). Consequently, although naïve PSCs consume less oxygen than MEFs and HL-1 cells, and consume more oxygen than EpiLCs and NSCs for ATP generation when normalized to cell number, naïve ESC mitochondria consume significantly more oxygen for ATP generation than mitochondria in primed EpiLCs, somatic stem cells (NSCs), somatic fibroblasts and HL-1 cardiomyocytes, when normalized to mitochondrial volume (Figures 1F, 1G, S1A, S1C, S1J and S1K). Mitochondria in naïve ESCs show the largest OXPHOS-ATP generation capacity among the tested cell lines when normalized to mitochondrial protein mass (Figures 1H and S1L). These results reveal the functionality of mitochondria in PSCs at distinct pluripotent states and somatic cells at different developmental stages, and partially explain the paradoxical opinions derived from the existing data in the literature. Our data support the hypothesis that naïve PSC mitochondria may be more active than mitochondria from PSC progeny when normalized to mitochondrial volume or mass.

The concomitant measurement of the oligomycin-sensitive oxygen consumption rate and the glycolytic rate, which represent OXPHOS-derived ATP production and glycolysis-derived ATP production respectively, enabled us to calculate the differential contributions of OXPHOS and glycolysis to ATP generation inside ESCs. Previous studies have shown that hPSCs, which are the equivalent of mouse primed PSCs, possess a significantly higher glycolysis rate than somatic fibroblasts ^29, 31^. Meanwhile, the glycolysis rate in naïve mouse PSCs is lower than that in primed PSCs, but significantly higher than that in somatic cells ^3, 4, 7, 8, 10, 23, 37^. Consistent with these findings, we found that mouse naïve ESCs have a higher rate of glycolysis than all the somatic cell lines tested here, and possess a lower glycolysis rate than primed EpiLCs (Figures S1B). However, in contrast with the existing proposition that ESCs favor glycolysis over OXPHOS for their energy supply, we established that OXPHOS accounts for ∼65% of total cellular ATP generation in naïve ESCs and ∼51% of total cellular ATP generation in primed ESCs (Figures 1E and S1C). These data led us to conclude that OXPHOS rather than glycolysis generates the majority of the cellular ATP in naïve ESCs, while OXPHOS generates a similar level of ATP as glycolysis in primed EpiLCs. These results provide objective views of the bioenergetics profile inside individual cells and highlight the critical role of OXPHOS for ATP generation in mESCs.

It has been demonstrated that naïve PSCs either transition into a diapause state or a primed differentiation stage upon signal stimulation, and both the diapause state and primed state show dramatic decreases of mitochondrial activity ^11, 38–40^. This raises a interesting question: could OXPHOS inhibition induce ESCs into the diapause state or primed state? Both diffusion map and scatterplots analyses indicate that OXPHOS inhibition induce ESCs into a unique state that is different from either the diapause or primed state (Figure S8). OXPHOS-inhibited ESCs appear closer to the diapause state than the primed state. However, OXPHOS-inhibited ESCs show significant differences with the diapause state, such as an inconsistent expression pattern of pluripotency genes etc. (Figure S8) ^38, 39^.

Interestingly, mouse naïve PSCs show a decreased glycolysis rate but an increased mitochondrial respiration compared to primed PSCs (Figures S1A and S1B) ^10^. However human naïve ESCs show an increase in both glycolysis rate and OXPHOS capacity compared to primed human ESCs ^9, 11^. Interesting areas for future investigation include the relative contribution of OXPHOS and glycolysis to ATP generation in human naïve and primed ESCs, and the relative bioenergetics requirement of human naïve ESCs compared to primed ESCs.

Mounting evidence suggests that the metabolite UDP-GlcNAc, the end product of the hexosamine biosynthesis pathway (HBP), plays critical roles in transcription, protein stability, metabolism, transport, *etc.* ^26, 41–43^. Our integrated transcriptome and metabolome analysis revealed that OXPHOS inhibition causes incomplete catabolism of glucose and abnormal metabolism of nucleotides, glutamine and acetyl-CoA, thus significantly decreasing the cellular UDP-GlcNAc level in ESCs (Figure S9). Accordingly, the O-GlcNAcylation and expression of pluripotency proteins significantly decreased independently of OGT expression, leading to disruption of pluripotency (Figure 4). O-GlcNAcylation was recently reported to have another role in pluripotency regulation, namely that O-GlcNAcylation of adenosylhomocysteinase (AHCY) promotes mESC pluripotency by maintaining a high level of H3K4me3 ^44^. It will be interesting to investigate whether the decrease of UDP-GlcNAc upon OXPHOS inhibition leads to compromised AHCY O-GlcNAcylation and thus affects ESC pluripotency. Together, these data suggest that the cellular UDP-GlcNAc concentration is tightly regulated by OXPHOS, and that UDP-GlcNAc serves as a critical cell fate regulator in PSCs.

The fact that reduced expression of the pluripotency genes Oct4 and Sox2 in response to OXPHOS inhibition lags behind their protein O-GlcNAcylation indicates that O-GlcNAcylation acts upstream of cell fate determination. These evidence led to the proposition to manipulate cell fate through metabolites. In support of this idea, exogenous supplementation of L-Pro induces mouse ESC mesenchymal transition ^45^, while deprivation of methionine induces human ESC differentiation ^46^. Interestingly, supplementation of α-ketoglutarate (αKG) to the culture media of naïve mESCs enhances self-renewal and inhibits differentiation, while adding α-ketoglutarate to cultures of human primed PSCs and mouse epiblast stem cells (EpiSCs) promotes differentiation, indicating the state-dependent effects of αKG in maintaining pluripotency or promoting differentiation^47, 48^. Taken together, these data suggest that despite being regulated by upstream signaling, metabolites may reciprocally exert its effects on signal transduction pathways that regulate downstream cell-fate determination, which points to the potential alternative strategy for exploiting metabolite manipulation to instruct directed differentiation.

In conclusion, the current study demonstrates that PSC mitochondria are in a super-active state and OXPHOS produces the majority of cellular ATP, which challenges the traditional concept that ESCs rely on glycolysis as their major source of energy. In addition, this study uncovered a previously unknown mechanism in which OXPHOS couples with the HBP pathway for UDP-GlcNAc generation to regulate pluripotency in mouse ESCs (Figures S10). The underlying mechanisms by which ESC mitochondria maintain a high respiration rate and favor UDP-GlcNAc production for pluripotency regulation remain to be further investigated.

## STAR METHODS

### KEY RESOURCES TABLE

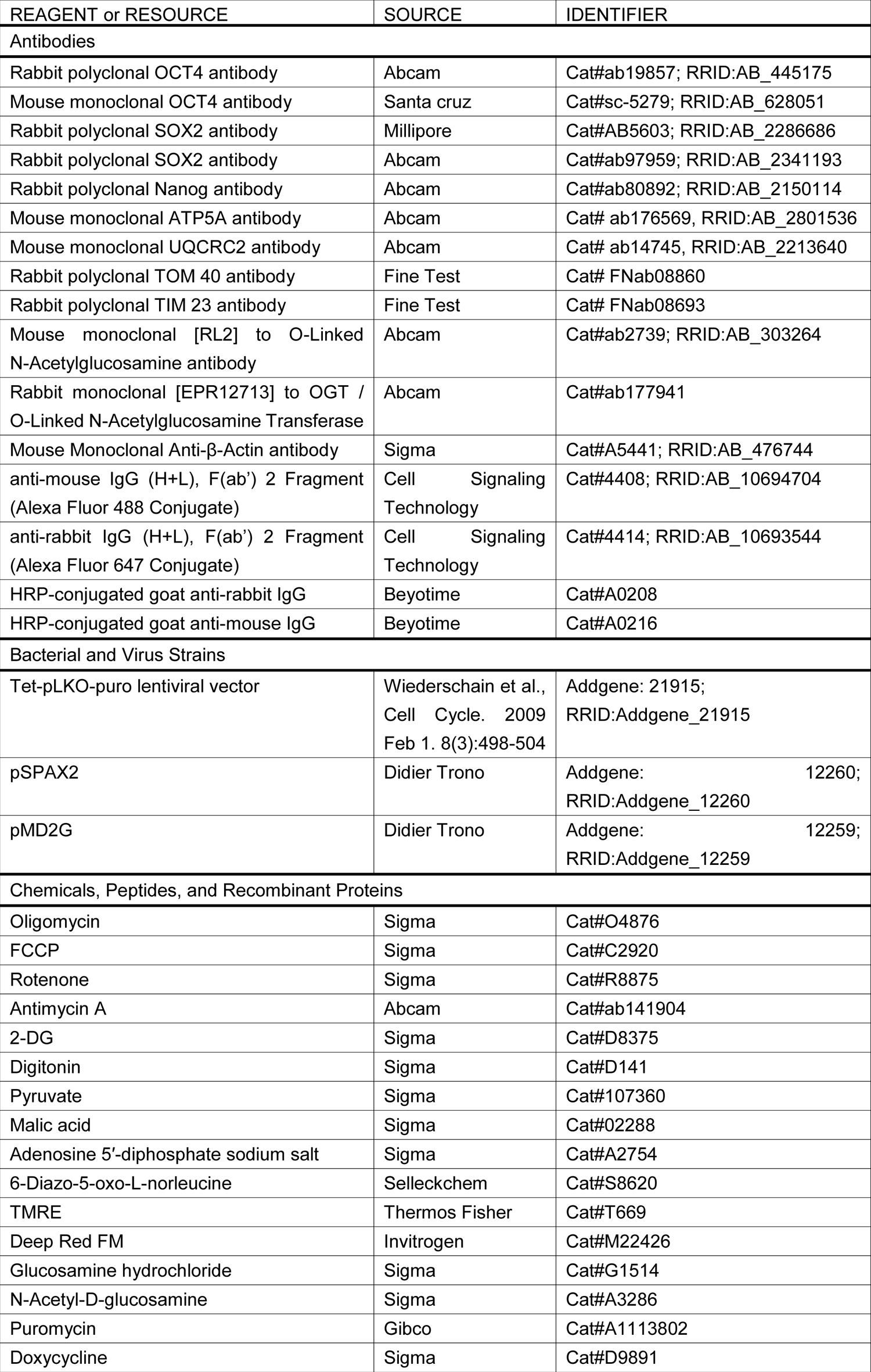
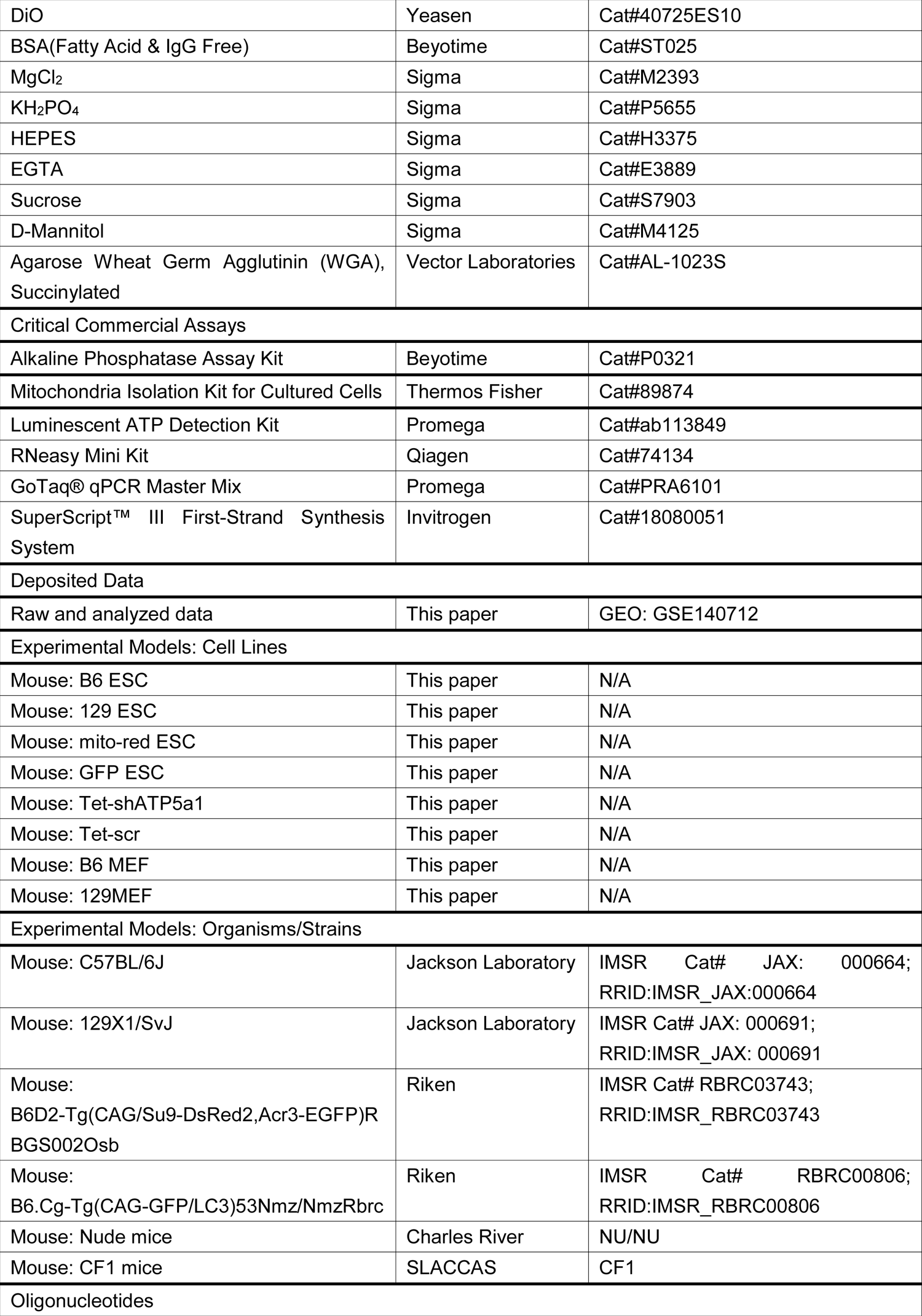
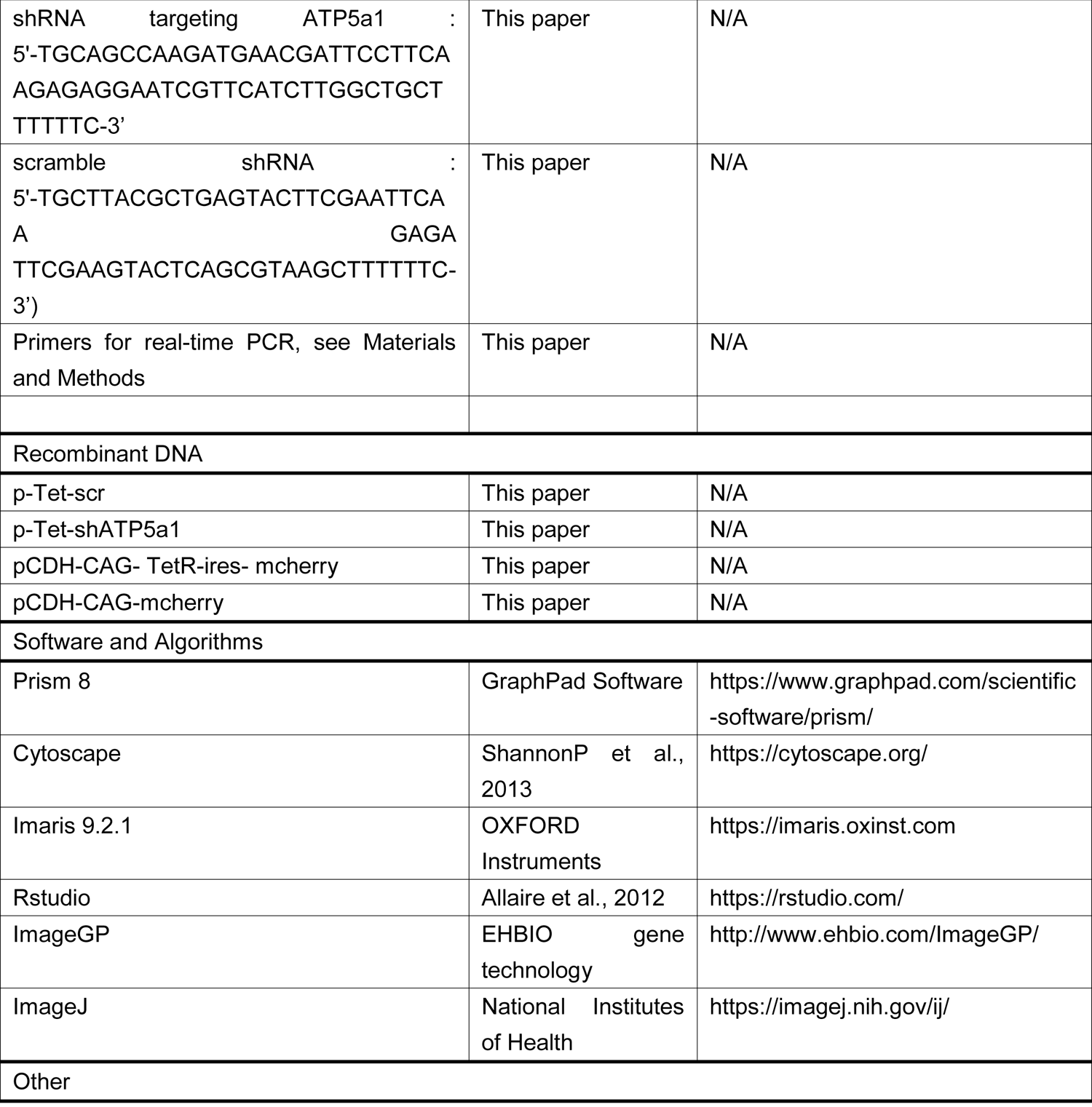

### LEAD CONTACT AND MATERIALS AVAILABILITY

Further detailed information and requests for resources and reagents should be directed to and will be fulfilled by the Lead Contact Tongbiao Zhao

### EXPERIMENTAL MODEL AND SUBJECT DETAILS

#### Reagents and Antibodies

Reagents: Oligomycin (Sigma, O4876), FCCP (Sigma, C2920), rotenone (Sigma, R8875), antimycin A (Abcam, ab141904), 2-DG (Sigma, D8375), GlcNAc (Sigma, A3286), puromycin (Gibco, A1113802), doxycycline (Dox; Sigma, D9891), glucosamine (Sigma, G1514), 6-diazo-5-oxo-L-norleucine (Selleckchem, S8620), digitonin (Sigma, #D141), ADP (Sigma, A2754), pyruvate (Sigma, 107360), malic acid (Sigma, 02288), EGTA (Sigma, E3889), HEPES (Sigma, H3375), sucrose (Sigma, S7903), D-mannitol (Sigma, M4125), MgCl_2_ (Sigma, M2393), KH_2_PO_4_ (Sigma, P5655), and BSA (Fatty Acid & IgG Free) (Beyotime, ST025) were used for cell treatments and Seahorse XF24 extracellular flux analysis. For extended resolution structured illumination imaging (SIM), DiO (Yeasen, 40725ES10) was used to stain cell membranes, Deep Red FM (Invitrogen, M22426) was used to stain mitochondria in EpiLCs and HL-1 cells, and TMRE (Thermos Fisher, T669) was used to stain mitochondria in NSCs. Mitochondria in ESCs and MEFs were labeled by mCherry.

Antibodies: anti-OCT4 (Abcam, ab19857, 1:1000 for western blot (WB); 1:100 for immunofluorescence analysis (IFA) of mESC), anti-OCT4 (Santa Cruz, sc-5279, 1:100 for IFA of blastocysts), anti-SOX2 (Millipore, AB5603, 1:100 for IFA of blastocysts), anti-SOX2 (Abcam, ab97959, 1:1000 for WB), anti-NANOG (abcam, ab80892, 1:1000 for WB), anti-O-linked N-acetylgucosamine (Abcam, ab2739, 1:1000 for WB, 1:50 for IFA of blastocysts), anti-OGT (Abcam, ab177941, 1:1000 for WB), anti-ATP5A (Abcam, ab176569, 1:1000 for WB), anti-UQCRC2 (Abcam, ab14745, 1:1000 for WB), anti-TOM 40 (Fine Test, FNab08860, 1:1000 for WB), anti-TIM 23 (Fine Test, FNab08693, 1:1000 for WB) and anti-β-ACTIN (A5441, Sigma, 1:1000 for WB) were used as primary antibodies in western blotting and immunofluorescence imaging assays; anti-mouse IgG (H+L), F(ab’)_2_ Fragment (Alexa Fluor 488 Conjugate) (CST, 4408, 1:1000) and anti-rabbit IgG (H+L), F(ab’)_2_ Fragment (Alexa Fluor 647 Conjugate)(CST, 4414, 1:1000) were used as secondary antibodies for immunofluorescence; HRP-conjugated goat anti-rabbit (A0208, Beyotime, 1:10000) and goat anti-mouse (A0216, Beyotime,1:10000) IgG were used as secondary antibodies for western blotting.

#### Plasmids and Lentivirus

The shRNA targeting ATP5a1 (5’-TGCAGCCAAGATGAACGATTCCTTCAAGAGAGGAATCGTTCATCTTGGCTGCTTTTTTC-3’) and the scramble shRNA (5’-TGCTTACGCTGAGTACTTCGAATTCAA GAGA TTCGAAGTACTCAGCGTAAGCTTTTTTC-3’) were cloned into the Tet-pLKO-puro lentiviral vector (a gift from Dmitri Wiederschain, Addgene, 21915), to create constructs named p-Tet-scr and p-Tet-shATP5a1 respectively. A cDNA encoding tetracycline repressor protein (TetR) was cloned into the pCDH-CAG-mcherry lentiviral expression vector, to create the construct named p-TetR. The lentivirus packaging vectors pSPAX2 and pMD2G were gifts from Didier Trono (Addgene, pSPAX, 12260; pMD2G, 12259). To prepare lentivirus, HEK293T cells were transduced with the plasmids p-Tet-scr/pSPAX2/pMD2G, p-Tet-shATP5a1/pSPAX2/pMD2G and p-TetR/pSPAX2/pMD2G using the calcium phosphate-DNA coprecipitation method. Medium containing the virus was collected 48 h after transfection.

#### Mice

All animal experiments were approved by the Ethics Committee in the Institute of Zoology, Chinese Academy of Sciences in accordance with the Guidelines for Care and Use of Laboratory Animals established by the Beijing Association for Laboratory Animal Science. C57BL/6 (Jackson Laboratory, 000664), 129X1/SvJ (Jackson Laboratory, 000691), mito-red B6D2-Tg(CAG/Su9-DsRed2,Acr3-EGFP)RBGS002Osb (Riken, RBRC03741) ^49^ and B6.Cg-Tg(CAG-GFP/LC3)53Nmz/NmzRbrc mice (Riken, RBRC00806) ^50^ were used for preparation of mouse embryonic fibroblasts (MEFs) and establishing ESC lines. C57BL/6 (Jackson Laboratory, 000664) mice were used for preparation of mouse neural stem cells (NSCs). Nude mice (Charles River, NU/NU) were used in teratoma formation assays. CF1 mice (SLACCAS) were used for feeder cell preparation and chimerism assays.

## METHOD DETAILS

### Generation of Stable ESC Lines

B6 ESCs were infected with lentivirus harboring TetR with the mcherry cassette, and the positive colonies were picked based on the mcherry fluorescence. Then the cells were infected with p-Tet-scr or p-Tet-shATP5a1, and selected with 1 μg/ml puromycin. The surviving colonies were picked and confirmed by sequencing before use in further experiments.

### Cell Culture

MEFs were isolated from embryonic day 13.5 (E13.5) fetal mice and passage 3 (P3) MEFs were used in each experiment. NSCs were isolated from cerebral cortex of E13.5 fetal mice. B6 ES (C57BL/6), J1 ES (129X1/SvJ) and Red-mito ES cell lines were isolated from E3.5 blastocysts and cultured on feeder layers for 5 d, and then routinely passaged. 1E iPS and 2E iPS cells were used as previously described ^51^. MEFs and HEK293T cells were maintained in DMEM High Glucose with 10% fetal bovine serum, 2 mM glutamine, 1 mM sodium pyruvate, and 1% penicillin/streptomycin at 37°C in 5% CO_2_. ESCs and iPSCs were maintained in KnockOut DMEM with 15% fetal bovine serum, 2 mM glutamine, 1 mM sodium pyruvate, 0.1 mM nonessential amino acids, 1% penicillin/streptomycin, 0.055 mM β-mercaptoethanol, and 1,000 U/ml leukemia inhibitory factor (Millipore). Specifically, the 2i ESCs were maintained in a 1:1 mix of DMEM/F12 (Gibco, 11302-33) and Neurobasal medium containing N2 supplements, B27 supplements, 10% KSR, 0.2% BSA, 1% penicillin/streptomycin, 0.1 mM β-mercaptoethanol, 2 mM glutamine and LIF supplemented with 1 μM PD03259010 (Stemgent), 3 μM CHIR99021 (Stemgent). The medium was changed every day. After three days of induction, the cells were passaged as cell clumps by dissociating with collagenase IV (1 mg/ml; Invitrogen), and maintained in the medium described as above but supplemented with 20% KSR. NSCs were maintained in Neurobasal medium containing B27 supplements, Glutamax, bFGF (20 ng/ml), EGF (20 ng/ml). HL-1 cells were maintained in MEM medium supplemented with 10% fetal bovine serum and 1% penicillin/streptomycin. EpiLCs were induced by plating 1.5 x 10^6^ ESCs on a 10 cm dish coated with human plasma fibronectin (16.7 mg/ml) in the 1:1 mix of DMEM/F12 (Gibco, 11302-33) and Neurobasal medium containing N2 supplements, B27 supplements, activin A (20 ng/ml), bFGF (12 ng/ml), and KSR (1%) ^52^. All cell culture reagents were purchased from Gibco unless indicated. For 20% or 50% inhibition of OXPHOS ATP production, ESCs were cultured in ESC medium containing 1 nM or 10 nM oligomycin respectively. For 20% or 50% inhibition of glycolysis ATP production, ESCs were cultured in ESC medium containing 5 mM or 10 mM 2-DG respectively. The established Tet-scr and Tet-shATP5a1 ESC lines were cultured in ESC medium containing 0, 125 ng/ml, or 250 ng/ml Dox. For HBP inhibition, ESCs were cultured in ESC medium containing 25 μM 6-diazo-5-oxo-L-norleucine. For the O-GlcNAcylation rescue assay, glucosamine (Sigma, G1514) or GlcNAc (Sigma, A3286) was supplemented as indicated.

### Immunofluorescence Microscopy

ESCs cultured on gelatin-coated glass slides were fixed with 4% paraformaldehyde for 30 min, washed with Dulbecco’s PBS (Hyclone), permeabilized by 0.2% Triton-X100 (Sigma Aldrich, X100) for 20 min, blocked with 2% bovine serum albumin (Sigma) for 1 h, stained with the indicated primary antibodies for 2 hour at 37°C, and then incubated with corresponding secondary antibodies for 2 h at room temperature.

For SIM, Mito-red ESCs and Mito-red MEFs cultured on gelatin-coated glass slides were fixed with 4% paraformaldehyde for 30 min, washed with Dulbecco’s PBS (Hyclone), and stained with 10 μM DiO for 5 min at 37°C; EpiLCs and HL-1 cells cultured on fibronectin-coated glass slides were stained with 20 nM Deep Red FM for 5 min at 37°C and then fixed with 4% paraformaldehyde for 30 min, washed with Dulbecco’s PBS (Hyclone), and stained with 10 μM DiO for 5 min at 37°C; NSCs cultured on Matrigel-coated glass slides were stained with 100 nM TMRE for 5 min at 37°C and stained with 10 μM DiO for 5 min at 37°C. Then the samples were visualized by a 3D-SIM built by Dr. Dong Li’s group ^20^. Briefly, the 3D-SIM was built based on an inverted fluorescence microscope (IX83, Olympus). The excitation light from a laser combiner equipped with 488 nm (500mW, Coherent Genesis Max), and 560 nm (500mW, MPB Communications, VFL-P-500-560) was passed through an acousto-optic tuneable filter (AOTF, AA Quanta Tech), which selected the excitation wavelength and controlled the power, coupled into an afterwards light path. The output beam from the AOTF was expanded to ∼15 mm and sent into a phase modulator consisting of a polarizing beam splitter, an achromatic half-wave plate (HWP; Bolder Vision Optik), and a ferroelectric spatial light modulator (SLM; Forth Dimension Displays, QXGA-3DM). The light diffracted by the grating pattern displayed on the SLM was directed to a polarization rotator (Meadowlark) to adjust the linear polarization of excitation light so as to maintain the s-polarization to maximize the interference pattern contrast. After selecting the 0 and ±1 diffraction orders, the excitation light was relayed into the back pupil of the objective (UPLSAPO100XS, Olympus), after which the collimated excitation light formed the interference pattern to illuminate the specimen. Meanwhile, the fluorescence was collected by the same objective, and relayed to a sCMOS camera (Hamamatsu, Orca Flash 4.0 v3) to acquire the raw images. The z-slices were sequentially acquired with 0.16 μm intervals to form a 3D-SIM image. The exposure time for each raw image was 10 ms at 20 W/cm.

The SIM images were rendered into 3D volumes and analyzed using Imaris V9.2.1. Cell surfaces were extracted using the Surface tools with manual editing, and the 3D structures of mitochondria were extracted by the Surface segmentation utility which detect objects based on local intensities. Images and videos were captured using the snapshot function and the animation function, respectively.

### Blastocysts

Two-cell embryos were isolated from E1.5 mouse embryos after the onset of pregnancy, and cultured in KSOM medium (Millipore, MR-121-D) at 37°C in 5% CO_2_ for 2 days. Then the blastocysts were randomly dived into 7 groups: Control (normally cultured in KSOM, 37°C, 5% CO_2_, 4 hours); 20%OI (cultured in KSOM supplemented with 1 nM oligomycin, 37°C, 5% CO_2_, 4 h); 20%OI+GlcNAc (cultured in KSOM supplemented with 1 nM oligomycin and 6 mM GlcNAc, 37°C, 5% CO_2_, 4 h), 20%GI (cultured in KSOM supplemented with 5 mM 2-DG, 37°C, 5% CO_2_, 4 h), 50%OI (cultured in KSOM supplemented with 10 nM oligomycin, 37°C, 5% CO_2_, 4 h); 50%OI+GlcNAc (cultured in KSOM supplemented with 10 nM oligomycin and 6 mM GlcNAc, 37°C, 5% CO_2_, 4 h) and 50%GI (cultured in KSOM supplemented with 10 mM 2-DG, 37°C, 5% CO_2_, 4 h).

### Inner Cell Mass Staining

After receiving the indicated treatments, *ex vivo* blastocysts were fixed with 4% paraformaldehyde for 20 min, washed with Dulbecco’s PBS (Hyclone), permeabilized by 0.5% Triton-X100 (Sigma Aldrich, X100) for 1 h, blocked with 2% bovine serum albumin (Sigma) for 1 h, washed with Dulbecco’s PBS (Hyclone), stained with the indicated primary antibodies overnight at 4°C and then incubated with corresponding secondary antibodies for 1 h at room temperature. Cell nuclei were counterstained with DAPI.

### XF24 Extracellular Flux Analysis

Respiration and acidification rates were measured in cells using a Seahorse XF24 analyzer (Seahorse Bioscience, North Billerica, MA). Briefly, ESCs, EpiLCs, NSCs, MEFs, or HL-1 cells were seeded on 0.1% gelatin-coated XF24 cell culture microplates (Seahorse Bioscience) at 5 × 10^4^ cells/well and incubated at 37°C in 5% CO_2_ overnight. For each XF24 extracellular flux assay, two microplates were prepared simultaneously. One microplate was used to measure the metabolism rates while the other one was used to determine the cell number at the time point when the measurement started.

The assays in Fig. 1E-H, Fig. S1A-C,H, Fig. S3A-C and Fig. S6A-B were performed in XF Base Medium (Agilent, 102353-100), supplemented with 25 mM glucose, 2 mM glutamine and 1 mM sodium pyruvate. Cells were washed twice and preincubated in this medium for 1 hour before testing. For ATP production determination, substrates and selective inhibitors were injected during the measurements to achieve final concentrations as following: port A: oligomycin (1 μM), port B: FCCP (0.3 μM), port C: rotenone (1 μM) and antimycin A (1 μM), port D: 2-DG (100 mM) (Fig.1. E-H, and Fig. S1A-C,H). To titrate the effects of oligomycin/2-DG on ATP production, inhibitors were injected during the measurements to achieve a series of final concentrations of oligomycin (1 nM, 5 nM, 10 nM, 100 nM, 200 nM, and 1000 μM), and 2-DG (2.5 mM, 5 mM, 10 mM and 100 mM) (Fig.S3A-C). For the mitochondrial stress test, inhibitors were injected to achieve final concentrations as following: port A: oligomycin (1 μM), port B: FCCP (0.3 μM), port C: rotenone (1 μM) and antimycin A (1 μM) (Fig.S6A-B). The oxygen consumption rate (OCR), extracellular acidification rate (ECAR) and protein production rate (PPR) values were further normalized to the number of cells present in each well.

The assays in Fig.S1I-L were performed in mannitol and sucrose (MAS) buffer (70 mM sucrose, 220 mM mannitol, 10 mM KH_2_PO_4_, 5 mM MgCl_2_, 2 mM HEPES and 1 mM EGTA, 4 mg/ml BSA; pH adjusted the to 7.2 with 0.1 M KOH). Substrates and selective inhibitors were injected during the measurements to achieve final concentrations as follows: port A: digitonin (25 μg/mL), pyruvate (5 mM), malate (2.5 mM) and ADP (1 mM); port B: oligomycin (1 μg/mL); port C: rotenone (1 μM) and antimycin A (20 μM) (Fig.1. E-H, and Fig. S1A-C, H) ^53^.

### ATP Production Calculation

The OCR, which drives mitochondrial ATP synthesis, can be calculated by addition of oligomycin (ATP synthase inhibitor), according to Equation 1 (E1) below. To convert the ATP-coupled OCR into the mitochondrial ATP production rate, the P/O ratios were estimated considering the stoichiometry of ATP phosphorylated per atom of oxygen consumed. Based on data from experiments on isolated mitochondria and considering the efficiency of the F1-F0 ATP synthase, the maximum theoretical P/O values obtained vary from 2.45 (palmitate oxidation) to 2.86 (full glycogen oxidation) (Brand, 2005; Mookerjee et al., 2017). A P/O ratio of 2.75 has been validated for calculating the OXPHOS ATP production rate in cells (Romero N et al Quantifying Cellular ATP Production Rate Using Agilent Seahorse XF Technology). Thus, the OXPHOS ATP production rate can be calculated according to Equation 2 (E2).

E1: OCR _ATP_ (pmol O_2_/min) = OCR _basal_ (pmol O_2_/min) – OCR _Oligo_ (pmol O_2_/min)

E2: OXPHOS ATP production rate (pmol ATP/min) = OCR _ATP_ (pmol O_2_/min) * 2 (pmol O/pmol O_2_) * 2.75 (P/O rate: pmol ATP/pmol O) Glycolysis converts glucose to lactic acid, accompanied by extrusion of one H^+^ (one proton) per lactate according to Equation 3 (E3) below. ECAR data are converted to PPR (proton production rate) based on the buffer capacity of the assay medium. The glycolytic ATP production rate is equivalent to the glycolytic proton production rate (PPR _glycolysis_), which is calculated as basal PPR minus non-glycolytic PPR (PPR _2DG_). Thus, the glycolytic ATP production rate can be calculated according to Equation 4 (E4).

E3: Glucose + 2 ADP + 2 Pi ➔ 2 Lactate + 2 ATP + 2 H_2_O + 2 H^+^

E4: Glycolytic ATP production rate (pmol ATP/min) = PPR _glycolysis_ (pmol H^+^/min) = PPR _basal_ (pmol H^+^/min) - PPR_2DG_ (pmol H^+^/min)

### Colony Formation Assay and AP Staining

ESCs were seeded at a density of 2000 cells/well in a six-well plate, and cultured for 7 days. The colonies were stained using an Alkaline Phosphatase Assay Kit (Beyotime, P0321), and the colony numbers were analyzed by Image-Pro Plus software.

### Mitochondria Isolation

The mitochondria from ESC, EpiLC, NSC, MEF, or HL-1 cell were isolated by a mitochondria isolation kit for cultured cells (Thermos Fisher, 89874).

### Western Blotting

Cells or the isolated mitochondria were lysed in RIPA buffer (50 mM Tris-HCl, pH 7.4, 150 mM NaCl, 0.5% sodium deoxycholate, 1% Nonidet P-40, 5 mM EGTA, 2 mM EDTA, 10 mM NaF) on ice for 30 min. A protease inhibitor cocktail tablet (04693116001, Roche) and 1 mM PMSF (ST506, Beyotime) were added during the lysis process. Cell lysates were subjected to SDS-PAGE and transferred to PVDF membranes (Millipore). Membranes were blocked with 5% nonfat milk for 1 h at room temperature and then probed with the indicated primary antibodies at 4°C overnight, followed by the appropriate HRP-conjugated secondary antibodies for 1 h at room temperature. After washing with TBST 3 times, the blotted membranes were visualized with chemiluminescent kits (Millipore). Densitometric analyses were performed using ImageJ software.

### Immunoprecipitation

The total protein of ESCs was extracted using a cell lysis buffer for Western and IP (Beyotime, P0013). Supernatants were isolated by centrifugation at 13000 × rpm for 15 min, and then incubated with sWGA agarose beads for 14 h at 4 °C with rotation (Vector laboratories, AL-1023S). The samples were centrifuged at 2500 × rpm for 5 min at 4 °C and washed five times with lysis buffer. The proteins coupled to beads were dissociated by boiling the samples at 100 °C for 10 min in 20 ul 5 × loading buffer (Beyotime, P0015L,) and then were subjected to western blot.

### Real-time PCR

Total RNA was extracted from samples with an RNeasy Mini Kit (Qiagen). About 1-2 μg total RNA was reverse-transcribed into cDNA using the SuperScript™ III First-Strand Synthesis System (Invitrogen). Qualitative PCR was performed with GoTaq® qPCR Master Mix (Promega) and a 7500 Fast Real-Time PCR System (Applied Biosystems). All samples were analyzed in duplicate and normalized to β-Actin. The primers used were as follows: Oct4: 5’-AGAGGATCACCTTGGGGTACA-3’ (forward), 5’-CGAAGCGACAGATGGTGGTC-3’ (reverse); Nanog: 5’-TCTTCCTGGTCCCCACAGTTT-3’ (forward), 5’-GCAAGAATAGTTCTCGGGATGAA-3’ (reverse); Sox2: 5’-GCGGAGTGGAAACTTTTGTCC-3’ (forward), 5’-CGGGAAGCGTGTACTTATCCTT-3’ (reverse); Esrrb: 5’-CAGGCAAGGATGACAGACG-3’ (forward), 5’-GAGACAGCACGAAGGACTGC-3’ (reverse); Rex1: 5’-CCCTCGACAGACTGACCCTAA-3’ (forward), 5’-TCGGGGCTAATCTCACTTTCAT-3’ (reverse); β-Actin: 5’-GGCTGTATTCCCCTCCATCG-3’ (forward), 5’-CCAGTTGGTAACAATGCCATGT-3’ (reverse).

### Teratoma Formation Assay

Approximately 2 × 10^6^ ESCs were subcutaneously injected into 10-week-old female nude mice. Four weeks later, teratomas were disassociated, weighed and fixed with 4% paraformaldehyde, embedded in paraffin, sectioned, and subjected to H&E staining.

### Chimerism Assay

After receiving the indicated treatments, the fluorescence-labeled ESCs were injected into blastocysts, then the injected blastocysts were transplanted into surrogate mice. The chimeric embryos (E13.5) were digested into single cells and subjected to FCAS analysis.

### Apoptosis Assay

The apoptosis assay was performed using an Alexa Fluor® 488 Annexin V/Dead Cell Apoptosis Kit (Invitrogen) according to manufacturer’s instructions. Briefly, cells were washed with PBS and re-suspended in binding buffer. The cells were incubated with annexin-V-FITC and propidium iodide (PI) for 20 min at RT, and then subsequently analyzed by flow cytometry. Cells that were annexin-V-negative and PI-negative were counted as viable cells.

### RNA-seq, Differential Gene Expression Analysis, and Function Enrichment Analysis

Total RNA was extracted from the indicated cell samples with an RNeasy Mini Kit (Qiagen). RNA sequencing was performed by Annoroad Gene Technology Corporation.

DESeq2 software was used for differential gene expression analysis. Briefly, the expression level of each gene per sample was estimated by DESeq2 according to linear regression, and the p-value was calculated with the Wald test. Finally, the p-value was corrected by the Benjamini-Hochberg (BH) method. Genes with q≤0.05 and |log2_ratio|≥1 were identified as differentially expressed genes (DEGs). The genome-wide heatmaps were plotted with GraphPad Prism 8 software.

GO (Gene Ontology, http://geneontology.org/) enrichment analysis of DEGs was implemented by the hypergeometric test, in which the p-value is calculated and adjusted according to the false discovery rate to give a q-value, and genes in the whole genome are used as the background data. GO terms with q<0.05 were considered to be significantly enriched. GO enrichment analysis assigns biological functions to DEGs. The GO enrichment map was plotted with imageGP.

Diffusion map is a nonlinear dimensionality reduction method where DC represents the diffusion coefficient ^54^. The correlation coefficient for omics data between two groups is calculated as R. All analyses were performed using two-tailed Student’s *t*-test (unless otherwise specified).

### Metabolomics Analysis

The ESCs were cultured in 10 cm dishes under feeder-free conditions (∼10^7 cells/dish) and treated as indicated for 48 h. The cells were washed with cold PBS 3 times and then were incubated in 2 ml 80% (vol/vol) methanol (pre-chilled to −80 °C) at −80°C for 1 h. Cell lysate/methanol mixtures were acquired by scraping, then centrifuged at 14000g for 20 min at 4°C. The protein content of the cell pellets was quantified for normalization. The metabolite-containing supernatants were removed to a new 1.5 ml tube and then dried to a pellet in a Speedvac. The metabolite pellet samples were submitted for the following metabolomic analysis. The targeted metabolomic experiment was analyzed by a TSQ Quantiva triple-stage quadrupole mass spectrometer (Thermo, CA). C18-based reverse-phase chromatography was utilized with 10 mM tributylamine, 15 mM acetate in water and 100% methanol as mobile phases A and B respectively. This analysis focused on TCA cycle, glycolysis pathway, pentose phosphate pathway, amino acid metabolism and purine metabolism. A 25 min gradient from 5% to 90% of mobile phase B was used. Positive-negative ion switching mode was performed for data acquisition. The resolution for Q1 and Q3 are both 0.7 FWHM. The source voltage was 3500 V for positive and 2500 V for negative ion mode. The source parameters are as follows: spray voltage, 3000 V; capillary temperature, 320 °C; heater temperature, 300 °C; sheath gas flow rate, 35; auxiliary gas flow rate, 10. Metabolite identification was based on a Tracefinder search with a home-built database containing about 300 compounds. Rstudio was used for differential metabolomics compounds analysis, and p-values were calculated with the two-tailed distribution *t*-test based on the two-sample isovariance hypothesis.

### Integrated Metabolome and Transcriptome Network Analysis

Based on the metabolites and transcripts detected in our experimental setting and the KEGG REACTION, KEGG ENZYME, KEGG COMPOUND, and KEGG GLYCAN databases, we integrated the metabolites and corresponding enzymes into a global network as previously described (Jha et al., 2015; Kanehisa et al., 2012). The network was annotated and plotted by Cytoscape software.

## STATISTICAL ANALYSIS

Statistical details and number of replicates are shown in the corresponding figure legends. Analyses were performed using SPSS Software. Statistical significance was calculated using unpaired *t* tests between the indicated groups, P-values <0.05 were considered as statistically significant. P-values are indicated by asterisks as follows: *P<0.05, **P<0.01, ***P<0.001.

## DATA AND CODE AVAILABILITY

The accession numbers for the raw data FASTQ files for the RNA sequencing data deposited in NCBI GEO is GEO: GSE140712. All relevant data supporting the findings of this study are also available from the lead contact upon request.

## SUPPLEMENTAL INFORMATION

Figure S1 – S10

Movie S1

Movie S2

Movie S3

Movie S4

Movie S5

Graphical Abstract

## Supporting information

Video S1-ESC SIM Image

Video S2-EpiLC SIM Image

Video S3-NSC SIM Image

Video S4-MEF SIM Image

Video S5-HL1 SIM Image

Table S3 Key resources table

Table S1 Trancriptomics analysis

Table S2 Metabolomics analysis

## ACKNOWLEDGEMENTS

We thank Dr. Dangsheng Li from *Cell Research*, Dr. Shyh-Chang, N. from IOZ-CAS, Dr. Jing Yang at Peking University and Drs. Suneng Fu and Maojun Yang from Tsinghua University for their constructive discussions and suggestions. We thank Dr Masaru Okabe at Osaka University Japan for providing the B6D2-Tg (CAG/Su9-DsRed2,Acr3-EGFP) RBGS002Osb mice (mito-red). We thank Dr Noboru Mizushima at the National Institute for Basic Biology Japan for providing the B6.Cg-Tg (CAG-GFP/LC3) 53Nmz/NmzRbrc mice. We thank Yun Feng from the Center for Biological Imaging (CBI), Institute of Biophysics, Chinese Academy of Science for help with SIM image analysis. This work was supported by grants from the National Key R&D Program of China 2018YFA0108402, the Strategic Priority Research Program of the Chinese Academy of Sciences XDA16030302, the Strategic Collaborative Research Program of the Ferring Institute of Reproductive Medicine Grant No.33, and the National Natural Science Foundation of China Program (31720103907, 31570995, 31621004), to T.Z. and (31400831) to J.C.

## AUTHOR CONTRIBUTIONS

J.C. and T.Z. designed the experiments, analyzed the data, and wrote the paper; J.C., M.L., K.L., X.S., N.S., Y.Y., X.W., Q.Z., L.W., X.C., L.Z. and X.L. performed the experiments under the overall coordination of T.Z.; C.L. and D.L. contributed to the measurement of the volume of mitochondria and cells; M.L., S.T. and Z.C. analyzed transcriptome and metabolomics data.

---

**Figure S1. Determination of the Contribution of Oxidative Phosphorylation and Glycolysis to ATP Generation in Mouse ESCs, EpiLCs, NSCs, MEFs and HL-1 cells** (A) Left, Oxygen consumption rate (OCR) measured by Seahorse Extracellular Flux Assay; right, oligomycin-sensitive oxygen consumption rates (Oligomycin OCR) in naïve ESCs, EpiLCs, NSCs, MEFs and HL-1 cells. Results are shown as mean ± SD of three independent experiments; ^**^P < 0.01; ^***^P < 0.001; Student’s *t*-test. (B) Left, Extracellular acidification rate (ECAR) measured by Seahorse Extracellular Flux Assay; right, glycolytic rate (ECAR) in naïve ESCs, EpiLCs, NSCs, MEFs and HL-1 cells. Results are shown as mean ± SD of three independent experiments; ^*^P < 0.05; ^**^P < 0.01; ^***^P < 0.001; Student’s *t*-test. (C) ATP production by oxidative phosphorylation (OXPHOS-ATP) or glycolysis (Glycolysis-ATP) in 10000 naïve ESCs, EpiLCs, NSCs, MEFs and HL-1 cells. The oligomycin-sensitive oxygen consumption rate is converted into the OXPHOS ATP production rate using a P/O ratio of 2.75, and ATP production by glycolysis is determined by a one-to-one relationship between the generation of protons and ATP. The ratio of proton production rate to ECAR (PPR/ECAR) was determined as 5.198 using the Seahorse XF24 analyzer based on the buffer capacity of the assay medium. Results are shown as mean ± SD of three independent experiments. n=3; ^***^P < 0.001; ns, not significant; Student’s *t*-test. (D) The level of UQCRC2 protein is constant in mitochondria from naïve ESCs, EpiLCs, NSCs, MEFs and HL-1 cells. The data are normalized to equal mitochondrial protein mass. Left, the level of TOM40, TIM23, ATP5A and UQCRC2 were detected by western blotting in 15 µg mitochondrial proteins isolated from naïve ESCs, EpiLCs, NSCs, MEFs and HL-1 cells. Right, results are shown as mean ± SD of three independent experiments. n=3; ^***^P < 0.001; ns, not significant; Student’s *t*-test. (E) Titration of cellular protein quantity in naïve ESCs, EpiLCs, NSCs, MEFs and HL-1 cells. (F) Absolute protein quantity in an individual naïve ESC, EpiLC, NSC, MEF and HL-1 cell. (G) The relative levels of UQCRC2 in an equal mass of total cellular protein. Left, the expression levels of UQCRC2, OCT4, NANOG, NESTIN and ACTIN in 15 µg cellular proteins from naïve ESCs, EpiLCs, NSCs, MEFs and HL-1 cardiomyocytes were detected by western blotting; right, results are shown as mean ± SD of three independent experiments. n=3; ^*^P < 0.05; Student’s *t*-test. (H) ATP production by OXPHOS or glycolysis in naïve ESCs, EpiLCs, NSCs, MEFs and HL-1 cells. Data are normalized to an equal mass of cellular protein. Results are shown as mean ± SD of three independent experiments. n=3; ^***^P < 0.001; ns, not significant; Student’s *t*-test. (I) OCR was measured by Seahorse Extracellular Flux Assay using digitonin-permeabilized cells fed complex-specific substrates of the preceding complex in the electron transport chain. DIG, digitonin; Pyr, pyruvate; Mal, malic acid; ADP, Adenosine-5’-diphosphate; Oligo, oligomycin; Rot, rotenone. (J) State 3 OCR (OCR values measured after injection of DIG/Pyr/Mal/ADP) in naïve ESCs is significantly higher than in EpiLCs and NSCs, and lower than in MEFs and HL-1 cells, when the data are normalized to equal numbers of cells. Results are shown as mean ± SD of three independent experiments. n=3; ^**^P < 0.01; ^***^P < 0.001; Student’s *t*-test. (K) State 3 OCR in naïve ESC is significantly higher than that in EpiLCs, NSCs, MEFs or HL-1 cells when the data are normalized to equal mitochondrial volume. Results are shown as mean ± SD of three independent experiments. n=3; ^*^P < 0.05; ^**^P < 0.01; Student’s *t*-test. (L) Relative state 3 OCR in naïve ESCs is significantly higher than in EpiLCs, NSCs, MEFs and HL-1 cells when the data are normalized to equal mitochondrial protein mass. Results are shown as mean ± SD of three independent experiments. n=3; ^*^P < 0.05; ^***^P < 0.001; Student’s *t*-test.

**Figure S2. ATP generation by OXPHOS and Glycolysis in Different Somatic Cell and Pluripotent Stem Cell Lines** (**A**) Left, ATP production by OXPHOS and glycolysis in different ESC and iPSC lines cultured in ESC medium suppled with fetal bovine serum; Right, the relative contribution of OXPHOS and glycolysis to ATP production in each cell line. **(B)** Left, ATP production by OXPHOS and glycolysis in different ESC and iPSC lines cultured in 2i medium; Right, the relative contribution of OXPHOS and glycolysis to ATP production in each cell line. **(C)** The contribution of OXPHOS to ATP production is increased in ESCs cultured in 2i medium compared with ESCs cultured in traditional serum conditions. **(D)** Left, ATP production by OXPHOS and glycolysis in different MEF cell lines; Right, the relative contribution of OXPHOS and glycolysis to ATP production in each MEF cell line. Results are shown as mean ± SD of 4 replicates from one representative of three independent experiments. ^*^P < 0.05; ^**^P < 0.01; ^***^P < 0.001; NS, not significant; Student’s *t*-test.

**Figure S3. Effects of Oligomycin/2-DG on ATP Production and Survival of ESCs** (**A**) ATP generation by OXPHOS (OXPHOS-ATP) is inhibited by oligomycin in a dose-dependent manner. Oligomycin inhibits 20% and 50% of maximum OXPHOS-ATP generation (designated as 20%OI and 50%OI) at concentrations of 1 nM and 10 nM, respectively. The OXPHOS-ATP inhibition rates are calculated as follows: [(OCR_basal_ - OCR_certain oligomycin concentration_) / (OCR_basal_ - OCR_1000 nM oligo_)] x 100%. (**B**) ATP generation by glycolysis (glycolysis-ATP) is inhibited by 2-DG in a dose-dependent manner. 2-DG inhibits 20% and 50% of maximum glycolysis-ATP generation (designated as 20%GI and 50%GI) at concentrations of 5 mM and 10 mM, respectively. The glycolysis-ATP inhibition rates are calculated as follows: [(ECAR _basal_ - ECAR _certain 2-DG concentration_) / (ECAR _basal_ - ECAR_100 mM 2-DG_)] x 100%. **(C)** 20%OI and 50%GI have similar inhibition effects on total cellular ATP generation. 20%OI, 50%OI, 20%GI and 50%GI accounts for 15.7% ± 5.8%, 36.0% ± 6.2%, 7.7% ± 1.3%, and 15.8% ± 3.6% of total cellular ATP inhibition respectively. Results are shown as mean ± SD from one representative of three independent experiments. 20%OI, 50%OI, n=3; 20%GI, 50%GI, n=4; ns, not significant; Student’s *t*-test. (**D**) The total cellular ATP contents were significantly decreased upon inhibition of OXPHOS, inhibition of glycolysis or inhibition of both. The cellular ATP contents were determined by a Luminescent ATP Detection Kit. Results are shown as mean ± SD of 6 replicates from one representative of three independent experiments, n=6; ^***^P < 0.001; Student’s *t*-test. (**E**) Moderate inhibition of OXPHOS or glycolysis for 48 h does not affect ESC apoptosis. Treatment of ESCs with 2-DG at 100 mM (100%GI) induces ESC death, serving as a positive control. The percentage of surviving cells (Annexin V^-^/PI^-^) was calculated from flow cytometry data. Results are shown as mean ± SD of 3 replicates from one representative of two independent experiments. ^***^P < 0.001; Student’s *t*-test.

**Figure S4. Oligomycin Directly Inhibits Self-renewal of ESCs and EpiLCs** (**A**) OCT4 expression is decreased upon moderate inhibition of OXPHOS but not glycolysis. Immunofluorescence images showing expression of OCT4 (green) in mcherry-labeled ESCs subjected to 20%OI, 50%OI, 20%GI or 50%GI for 48 h. Bars, 50 µm. (**B**) Withdrawing oligomycin causes partial recovery of self-renewal in 20%OI and 50%OI ESCs in a time-dependent manner. The ESCs were seeded and subjected to 20%OI or 50%OI treatments for the indicated time, then the oligomycin was withdrawn. After a total culture period of 7 days, alkaline phosphatase staining was used to test the colony formation ability. Left, representative photos of ESC colonies stained by alkaline phosphatase; Right, statistical analysis of the number of alkaline phosphatase-positive colonies. Results are shown as mean ± SD of 3 independent experiments. ^***^ P < 0.001; Student’s *t*-test. (**C)** Images of representative teratomas formed by 20%GI and 50%GI ESCs. Bars, 1 cm. (**D**) Teratomas generated by control, 20%GI and 50%GI ESCs contain three embryonic germ layers. (**E**) Oligomycin inhibits self-renewal of EpiLCs dose-dependently. Left, representative photos of EpiLC colonies stained by alkaline phosphatase; right, statistical analysis of the number of alkaline phosphatase-positive colonies. Results are shown as mean ± SD of 3 replicates from one representative of 3 independent experiments. ^***^P < 0.001; Student’s *t*-test.

**Figure S5. Dox-inducible Knockdown of ATP Synthase** (**A**) The mRNA expression of ATP5a1 is inhibited by Dox in a dose-dependent manner. Results are shown as mean ± SD from 3 independent experiments. ^*^P < 0.05; ^***^P < 0.001; Student’s *t*-test. (**B**) Inhibition of ATP5a1 expression results in decreased expression of OCT4 (green). Bars, 50 μm.

**Figure S6. Genetic Inhibition of Oxidative Phosphorylation Causes Loss of ESC Identity** (**A**) Oxygen consumption rate (OCR) measurements on stable ESC lines expressing either scramble shRNA or shRNA targeting ATP5a1 under Dox control. (**B**) The ATP production-related OCR of Tet-shATP5a1 ESC decreases upon Dox treatment. Results are shown as mean ± SD of triplicates from one representative of three independent experiments. ^**^ P < 0.01; Student’s *t*-test. (**C**) Knockdown of ATP5a1 inhibits ESC self-renewal. Left, representative photos of ESC colonies stained with alkaline phosphatase. Right, statistical analysis of alkaline phosphatase-positive colonies. Results are shown as mean ± SD from 3 independent experiments. ^***^P < 0.001; Student’s *t*-test. (**D**) ATP5a1 knockdown decreases pluripotency gene expression. Results are shown as mean ± SD of 3 independent experiments. ^*^ P < 0.05; ^**^P < 0.01; ^***^P < 0.001; Student’s *t*-test. (**E**) Diagram of the chimeric mouse formation assay. (**F**) Knockdown of ATP5a1 decreases the ESC chimera rate. The mCherry-positive cells detected by FACS indicate the number of cells in each chimeric embryo that were derived from the originally injected cells. (**G**) Summary of data from chimeric embryos. Each dot represents the percentage of mCherry^+^ cells in an individual chimeric embryo. Scr, n=36; ATP5a1, n=30. ^***^P < 0.001; Student’s *t*-test.

**Figure S7. O-GlcNAcylation of Pluripotency Factors is Essential for ESC Identity** (**A**) Heatmap of expression of genes involved in the HBP pathway. (**B**) 50%OI treatment causes decreased O-GlcNAcylation and expression of SOX2 and OCT4 in naïve-state ESCs. (**C**) Statistical analysis of the western blot results shown in **B**. n=3; ^*^P < 0.05; ^**^P < 0.01; ^***^P < 0.001; ns, not significant; Student’s *t*-test. (**D**) Inhibition of the HBP pathway leads to breakdown of ESC identity. Left, HBP pathway inhibition by Don causes decreased expression of OCT4 and SOX2; right, representative images of ESC colonies stained with alkaline phosphatase. Don, 6-diazo-5-oxo-L-norleucine. (E) Both GlcNAc and glucosamine rescue the decreased O-GlcNAcylation and expression of OCT4 caused by OXPHOS inhibition. (**F**) Compromised expression of pluripotency genes led by OXPHOS inhibition is partially rescued by GlcNAc. Values displayed correspond to the expression level in the indicated sample scaled by the mean expression of each gene across samples. (**G**) OXPHOS inhibition causes altered expression of genes in the whole protein-coding transcriptome, and this effect is partially rescued by GlcNAc. Values displayed correspond to the expression level in the indicated sample scaled by the mean expression of each gene across samples. (**H**) Inhibition of OXPHOS rather than glycolysis reduces O-GlcNAcylation and expression of SOX2 in the inner cell mass of *ex vivo*-cultured blastocysts. These effects are ameliorated by adding GlcNAc. Confocal immunofluorescence microscopy was carried out using antibodies specific to O-GlcNAc (green) and SOX2 (red). The nuclei were stained with DAPI. Bars, 25 µm. (**I**) Statistical analysis of the immunofluorescence microscopy results shown in **H**. Control, n=10; 20%OI, n=11; 20%OI+GlcNAC, n=10; 20%GI, n=7; 50%OI, n=9; 50%OI+GlcNAC, n=9; 50%GI, n=10; ^*^P < 0.05; ^**^P < 0.01; ^***^P < 0.001; ns, not significant; Student’s *t*-test.

**Figure S8. OXPHOS Inhibition Induces ESCs into a State Which is Different to the Diapause- or Primed-State** (**A**) Diffusion map of ESCs, 20%OI- or 50%OI-treated ESCs and ESCs(2i/LIF), EPIs(E4.5), diapause EPIs or EPIs(E5.5) shows that OXPHOS inhibition induces ESCs into a unique state, which is closer to the diapause-state than to the primed-state (EPIs, epiblasts). (**B**) Scatterplots of gene expression levels that are significantly changed in the comparison of 20%OI/WT and Diapause/E4.5 (top left), 20%OI/WT and Primed(E5.5)/E4.5 (top right), 50%OI/WT and Diapause/E4.5 (bottom left) or 50%OI/WT and Primed(E5.5)/E4.5 (bottom right). The x axis shows the log_2_ fold change (log_2_ Fc) between 20%OI-(or 50%OI-) ESCs and control ESCs; the y axis shows the log_2_ Fc between diapause epiblasts (or E5.5 primed epiblasts) and E4.5 epiblasts. The dots represent the set of all genes. Differentially expressed genes (|FC|≥1.5 and p≤0.05) are shown as red dots. The numbers of DEGs (red) and total genes (gray) are labeled in every corresponding quadrant. The linear equation is fitted by all the dots in the four quadrants, and the slope of this linear fitting equation partially reflects the correlation of the x and y axes. R^2^ values display correlation coefficients. Data for ESCs(2i/LIF), EPIs(E4.5), diapause EPIs and EPIs(E5.5) are from Boroviak et al. 2015.

**Figure S9. Network Integration of Metabolome and Transcriptome Data Reveals that Inhibition of OXPHOS Leads to Defective UDP-GlcNAc Biosynthesis** A global combined network that connects metabolite nodes to reaction nodes was constructed based on the detected metabolites and the corresponding enzymes. The node of each reaction is connected to its respective substrates and products (solid circles). Enzymes that catalyze representative reactions are shown (diamonds). The relative levels of metabolites and enzymes in 20%OI ESCs compared to control ESCs is indicated by different colors (red, upregulated; green, downregulated; blue, not measured), and the significance is indicated by dot size (the bigger the more significant).

**Figure S10. UDP-GlcNAc Links Oxidative Phosphorylation to Pluripotency** ESC mitochondria have a high ATP generation capacity. Mitochondrial respiration produces the majority of cellular ATP in ESC, and couples with the hexosamine biosynthesis pathway to generate UDP-GlcNAc for O-GlcNAcylation of pluripotency factors like OCT4 and SOX2, thereby safeguarding ESC identity. Moderate inhibition of mitochondrial respiration leads to incomplete catabolism of glucose together with abnormal metabolism of glutamine, nucleotides, and acetyl-CoA, resulting in decreased UDP-GlcNAc generation and compromised pluripotency. This effect can be ameliorated by direct supplementation of GlcNAc.

