## supplemental figures for "Oxidative phosphorylation safeguards pluripotency via UDP-N-acetylglucosamine"

**Figure S1. Determination of the Contribution of Oxidative Phosphorylation and Glycolysis to ATP Generation in Mouse ESC, EpiLC, NSC, MEF and HL-1**

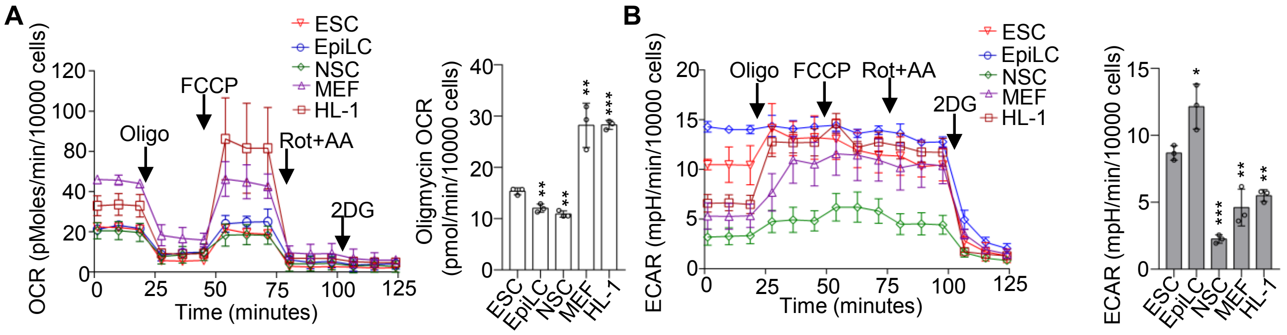


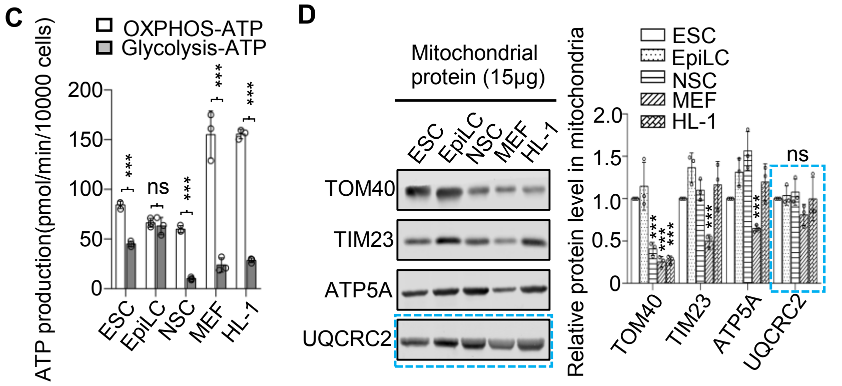


F

| Cell line | Protein(pg) /cell |
|---|---|
| ESC | 257.43±18.28 |
| EpiLC | 222.05±3.80 |
| NSC | 146.14±1.19 |
| MEF | 535.00±0.56 |
| HL-1 | 563.33±2.22 |

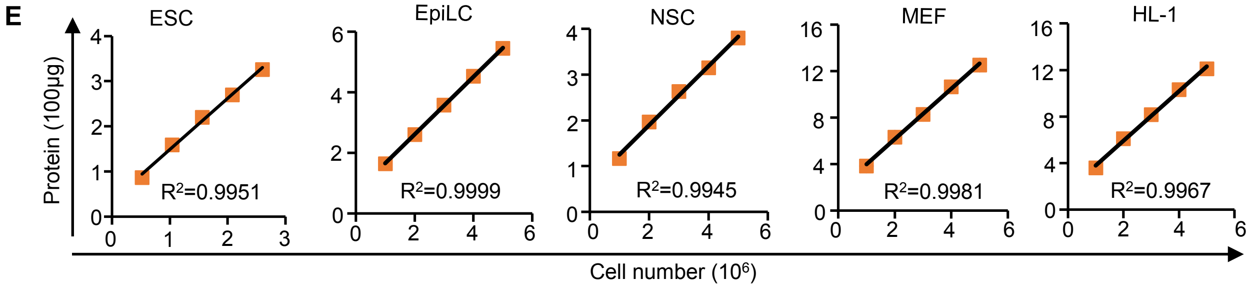


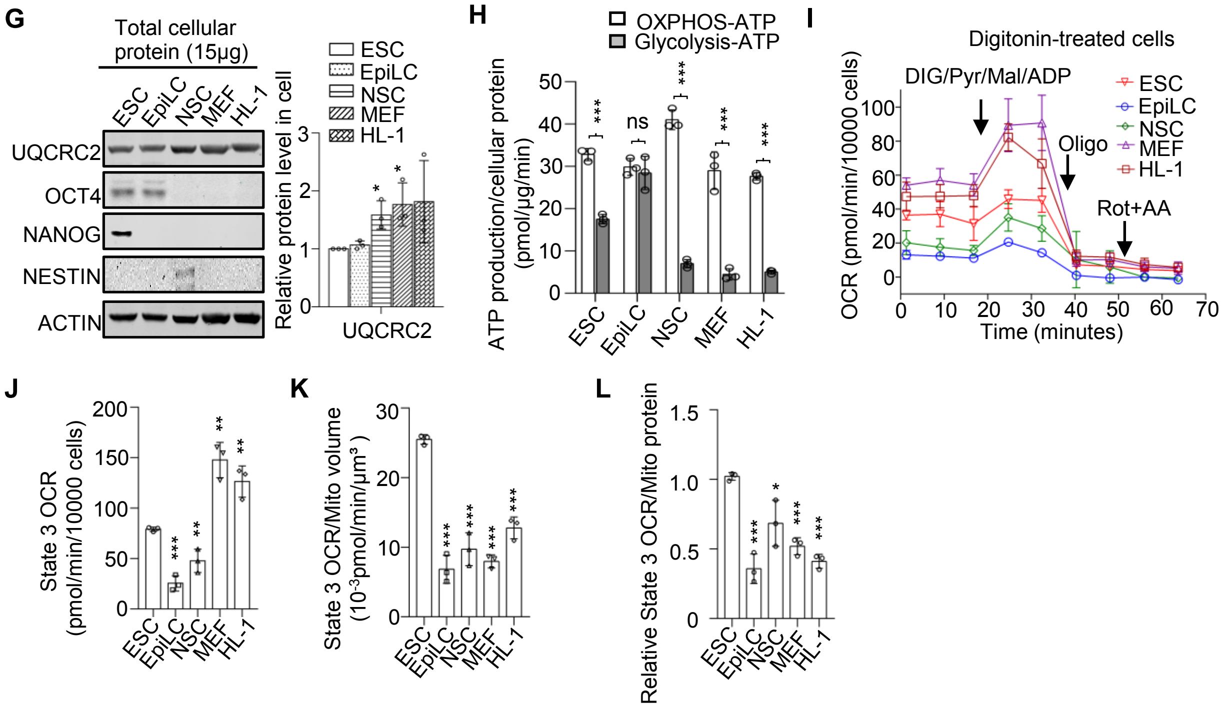

**Figure S1. Determination of the Contribution of Oxidative Phosphorylation and Glycolysis to ATP Generation in Mouse ESCs, EpiLCs, NSCs, MEFs and HL-1 cells** (**A**) Left, Oxygen consumption rate (OCR) measured by Seahorse Extracellular Flux Assay; right, oligomycin-sensitive oxygen consumption rates (Oligomycin OCR) in naïve ESCs, EpiLCs, NSCs, MEFs and HL-1 cells. Results are shown as mean ± SD of three independent experiments; **P < 0.01; ***P < 0.001; Student's *t*-test. (**B**) Left, Extracellular acidification rate (ECAR) measured by Seahorse Extracellular Flux Assay; right, glycolytic rate (ECAR) in naïve ESCs, EpiLCs, NSCs, MEFs and HL-1 cells. Results are shown as mean ± SD of three independent experiments; *P < 0.05; **P < 0.01; ***P < 0.001; Student's *t*-test. (**C**) ATP production by oxidative phosphorylation (OXPHOS-ATP) or glycolysis (Glycolysis-ATP) in 10000 naïve ESCs, EpiLCs, NSCs, MEFs and HL-1 cells. The oligomycin-sensitive oxygen consumption rate is converted into the OXPHOS ATP production rate using a P/O ratio of 2.75, and ATP production by glycolysis is determined by a one-to-one relationship between the generation of protons and ATP. The ratio of proton production rate to ECAR (PPR/ECAR) was determined as 5.198 using the Seahorse XF24 analyzer based on the buffer capacity of the assay medium. Results are shown as mean ± SD of three independent experiments. n=3; ***P < 0.001; ns, not significant; Student's *t*-test. (**D**) The level of UQCRC2 protein is constant in mitochondria from naïve ESCs, EpiLCs, NSCs, MEFs and HL-1 cells. The data are normalized to equal mitochondrial protein mass. Left, the level of TOM40, TIM23, ATP5A and UQCRC2 were detected by western blotting in 15 µg mitochondrial proteins isolated from naïve ESCs, EpiLCs, NSCs, MEFs and HL-1 cells. Right, results are shown as mean ± SD of three independent experiments. n=3; ***P < 0.001; ns, not significant; Student's *t*-test. (**E**) Titration of cellular protein quantity in naïve ESCs, EpiLCs, NSCs, MEFs and HL-1 cells. (**F**) Absolute protein quantity in an individual naïve ESC, EpiLC, NSC, MEF and HL-1 cell. (**G**) The relative levels of UQCRC2 in an equal mass of total cellular protein. Left, the expression levels of UQCRC2, OCT4, NANOG, NESTIN and ACTIN in 15 µg cellular proteins from naïve ESCs, EpiLCs, NSCs, MEFs and HL-1 cardiomyocytes were detected by western blotting; right, results are shown as mean ± SD of three independent experiments. n=3; *P < 0.05; Student's *t*-test. (**H**) ATP production by OXPHOS or glycolysis in naïve ESCs, EpiLCs, NSCs, MEFs and HL-1 cells. Data are normalized to an equal mass of cellular protein. Results are shown as mean ± SD of three independent experiments. n=3; ***P < 0.001; ns, not significant; Student's *t*-test. (**I**) OCR was measured by Seahorse Extracellular Flux Assay using digitonin-permeabilized cells fed complex-specific substrates of the preceding complex in the electron transport chain. DIG, digitonin; Pyr, pyruvate; Mal, malic acid; ADP, Adenosine-5'-diphosphate; Oligo, oligomycin; Rot, rotenone. (**J**) State 3 OCR (OCR values measured after injection of DIG/Pyr/Mal/ADP) in naïve ESCs is significantly higher than in EpiLCs and NSCs, and lower than in MEFs and HL-1 cells, when the data are normalized to equal numbers of cells. Results are shown as mean ± SD of three independent experiments. n=3; **P < 0.01; ***P < 0.001; Student's *t*-test. (**K**) State 3 OCR in naïve ESC is significantly higher than that in EpiLCs, NSCs, MEFs or HL-1 cells when the data are normalized to equal mitochondrial volume. Results are shown as mean ± SD of three independent experiments. n=3; *P < 0.05; **P < 0.01; Student's *t*-test. (**L**) Relative state 3 OCR in naïve ESCs is significantly higher than in EpiLCs, NSCs, MEFs and HL-1 cells when the data are normalized to equal mitochondrial protein mass. Results are shown as mean ± SD of three independent experiments. n=3; *P < 0.05; ***P < 0.001; Student's *t*-test.

**Figure S2. ATP generation by OXPHOS and Glycolysis in Different Somatic Cell and Pluripotent Stem Cell Lines**

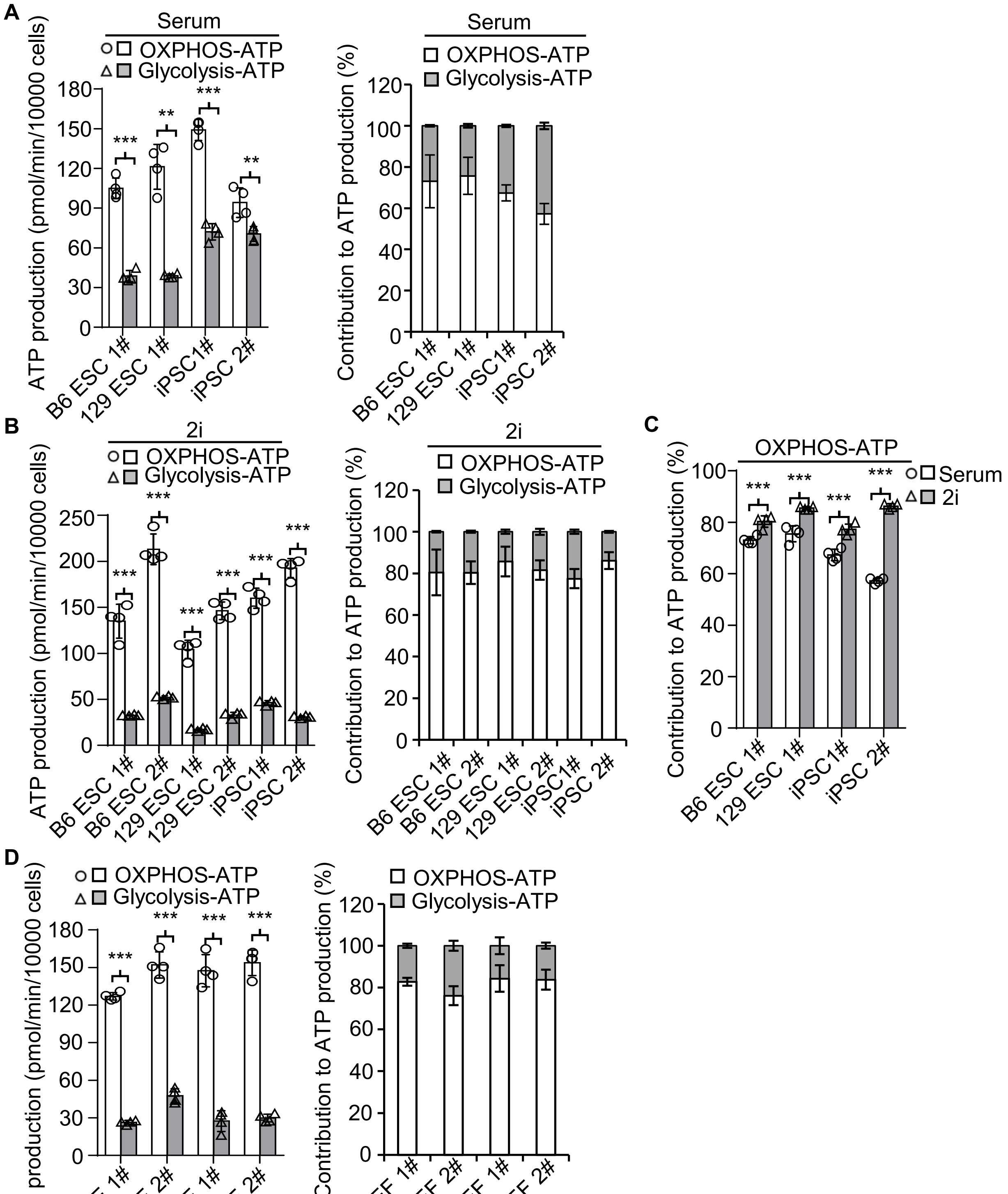


**Figure S2. ATP generation by OXPHOS and Glycolysis in Different Somatic Cell and Pluripotent Stem Cell Lines**

**(A)** Left, ATP production by OXPHOS and glycolysis in different ESC and iPSC lines cultured in ESC medium suppled with fetal bovine serum; Right, the relative contribution of OXPHOS and glycolysis to ATP production in each cell line. **(B)** Left, ATP production by OXPHOS and glycolysis in different ESC and iPSC lines cultured in 2i medium; Right, the relative contribution of OXPHOS and glycolysis to ATP production in each cell line. **(C)** The contribution of OXPHOS to ATP production is increased in ESCs cultured in 2i medium compared with ESCs cultured in traditional serum conditions. **(D)** Left, ATP production by OXPHOS and glycolysis in different MEF cell lines; Right, the relative contribution of OXPHOS and glycolysis to ATP production in each MEF cell line. Results are shown as mean ± SD of 4 replicates from one representative of three independent experiments. *P < 0.05; **P < 0.01; ***P < 0.001; NS, not significant; Student's *t*-test.

**Figure S3. Effects of Oligomycin/2-DG on ATP Production and Survival of ESCs**

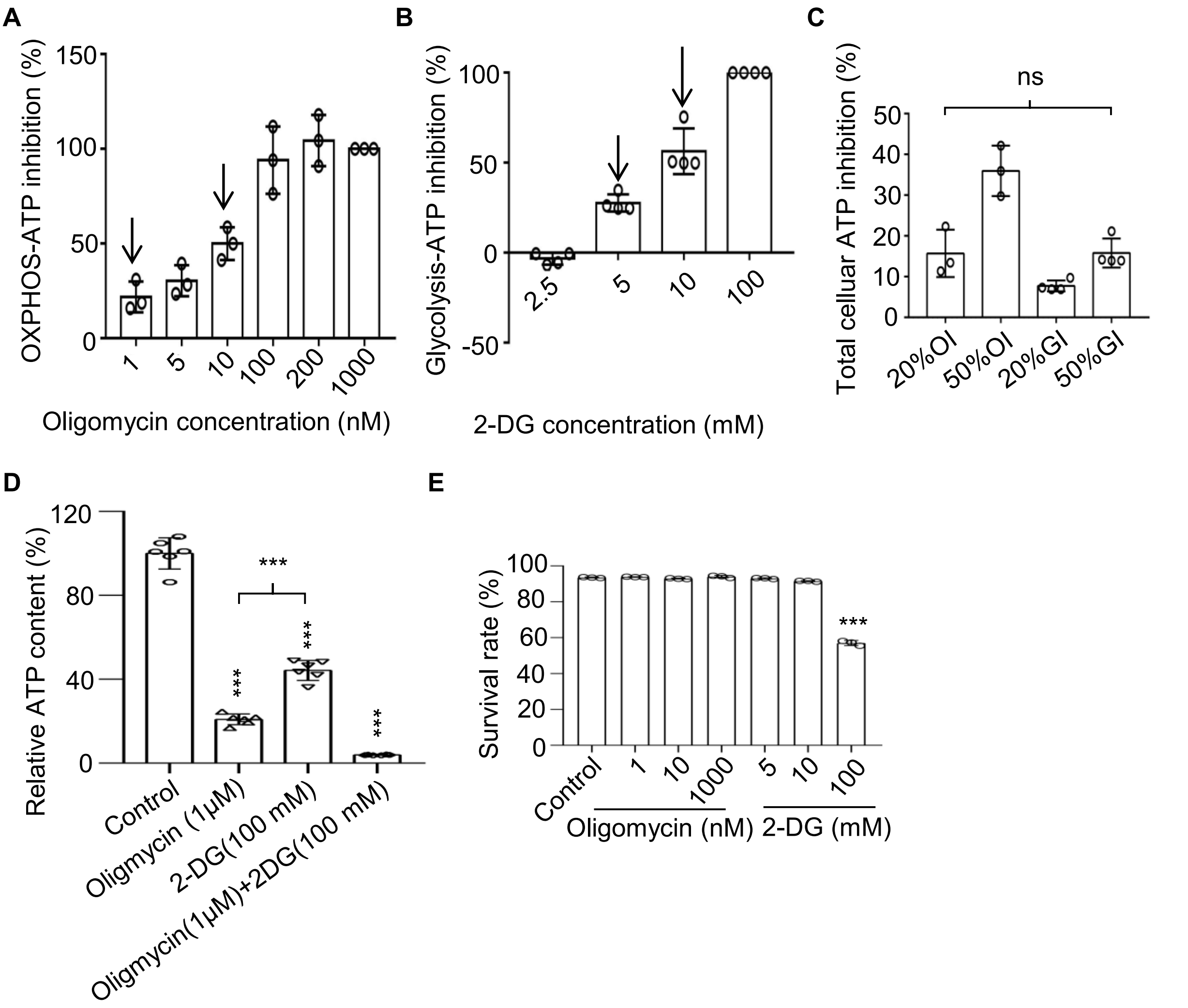


**Figure S3. Effects of Oligomycin/2-DG on ATP Production and Survival of ESCs** (**A**) ATP generation by OXPHOS (OXPHOS-ATP) is inhibited by oligomycin in a dose-dependent manner. Oligomycin inhibits 20% and 50% of maximum OXPHOS-ATP generation (designated as 20%OI and 50%OI) at concentrations of 1 nM and 10 nM, respectively. The OXPHOS-ATP inhibition rates are calculated as follows: [($OCR_{basal}$ - $OCR_{certain\ oligomycin\ concentration}$) / ($OCR_{basal}$ - $OCR_{1000\ nM\ oligo}$)] x 100%. (**B**) ATP generation by glycolysis (glycolysis-ATP) is inhibited by 2-DG in a dose-dependent manner. 2-DG inhibits 20% and 50% of maximum glycolysis-ATP generation (designated as 20%GI and 50%GI) at concentrations of 5 mM and 10 mM, respectively. The glycolysis-ATP inhibition rates are calculated as follows: [($ECAR_{basal}$ - $ECAR_{certain\ 2\text{-}DG\ concentration}$) / ($ECAR_{basal}$ - $ECAR_{100\ mM\ 2\text{-}DG}$)] x 100%. **(C)** 20%OI and 50%GI have similar inhibition effects on total cellular ATP generation. 20%OI, 50%OI, 20%GI and 50%GI accounts for 15.7% ± 5.8%, 36.0% ± 6.2%, 7.7% ± 1.3%, and 15.8% ± 3.6% of total cellular ATP inhibition respectively. Results are shown as mean ± SD from one representative of three independent experiments. 20%OI, 50%OI, n=3; 20%GI, 50%GI, n=4; ns, not significant; Student's *t*-test. (**D**) The total cellular ATP contents were significantly decreased upon inhibition of OXPHOS, inhibition of glycolysis or inhibition of both. The cellular ATP contents were determined by a Luminescent ATP Detection Kit. Results are shown as mean ± SD of 6 replicates from one representative of three independent experiments, n=6; ***P < 0.001; Student's *t*-test. (**E**) Moderate inhibition of OXPHOS or glycolysis for 48 h does not affect ESC apoptosis. Treatment of ESCs with 2-DG at 100 mM (100%GI) induces ESC death, serving as a positive control. The percentage of surviving cells (Annexin $V^-$/$PI^-$) was calculated from flow cytometry data. Results are shown as mean ± SD of 3 replicates from one representative of two independent experiments. ***P < 0.001; Student's *t*-test.

**Figure S4. Oligomycin Directly Inhibits Self-renewal of ESC and EpiLC**

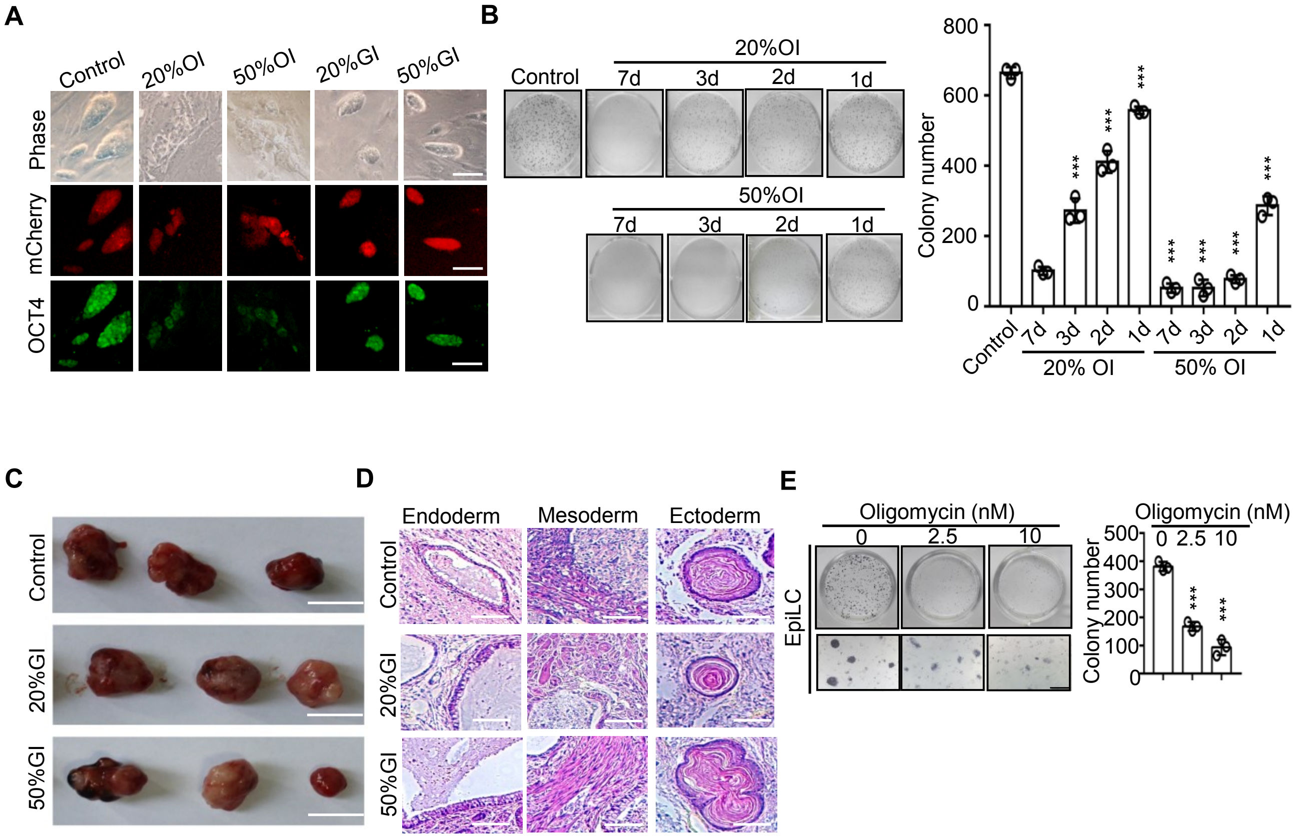


**Figure S4. Oligomycin Directly Inhibits Self-renewal of ESCs and EpiLCs** (**A**) OCT4 expression is decreased upon moderate inhibition of OXPHOS but not glycolysis. Immunofluorescence images showing expression of OCT4 (green) in mcherry-labeled ESCs subjected to 20%OI, 50%OI, 20%GI or 50%GI for 48 h. Bars, 50 μm. (**B**) Withdrawing oligomycin causes partial recovery of self-renewal in 20%OI and 50%OI ESCs in a time-dependent manner. The ESCs were seeded and subjected to 20%OI or 50%OI treatments for the indicated time, then the oligomycin was withdrawn. After a total culture period of 7 days, alkaline phosphatase staining was used to test the colony formation ability. Left, representative photos of ESC colonies stained by alkaline phosphatase; Right, statistical analysis of the number of alkaline phosphatase-positive colonies. Results are shown as mean ± SD of 3 independent experiments. ***P < 0.001; Student's *t*-test. (**C)** Images of representative teratomas formed by 20%GI and 50%GI ESCs. Bars, 1 cm. (**D**) Teratomas generated by control, 20%GI and 50%GI ESCs contain three embryonic germ layers. (**E**) Oligomycin inhibits self-renewal of EpiLCs dose-dependently. Left, representative photos of EpiLC colonies stained by alkaline phosphatase; right, statistical analysis of the number of alkaline phosphatase-positive colonies. Results are shown as mean ± SD of 3 replicates from one representative of 3 independent experiments. ***P < 0.001; Student's *t*-test.

**Figure S5. Dox-inducible Knockdown of ATP Synthase**

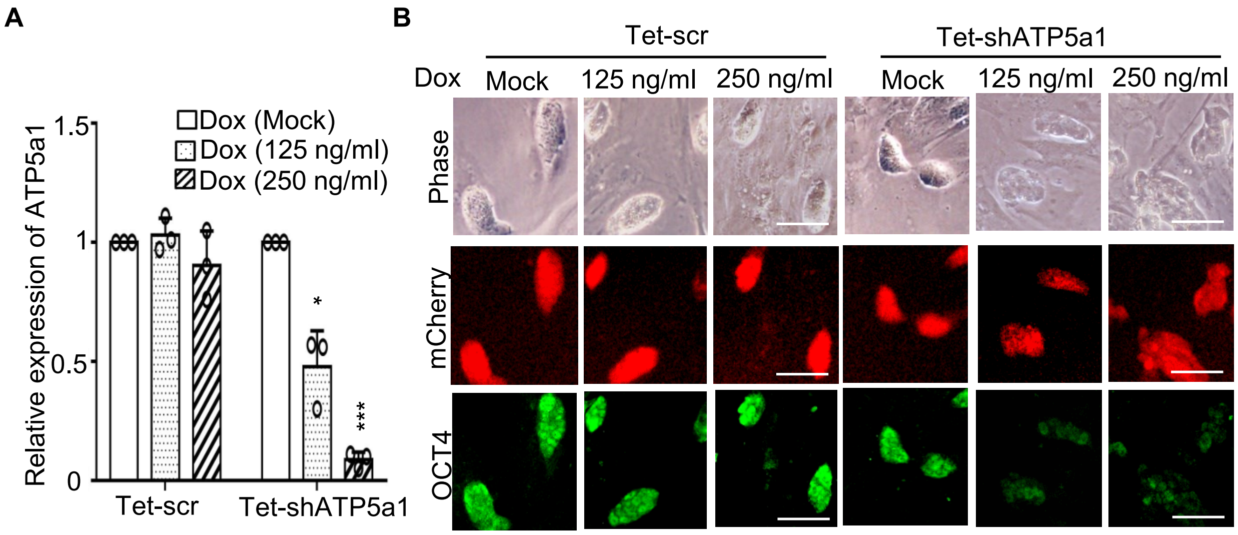


**Figure S5. Dox-inducible Knockdown of ATP Synthase** (**A**) The mRNA expression of ATP5a1 is inhibited by Dox in a dose-dependent manner. Results are shown as mean ± SD from 3 independent experiments. *P < 0.05; ***P < 0.001; Student's *t*-test. (**B**) Inhibition of ATP5a1 expression results in decreased expression of OCT4 (green). Bars, 50 μm.

### Figure S6. Genetic Inhibition of Oxidative Phosphorylation Causes Loss of ESC Identity

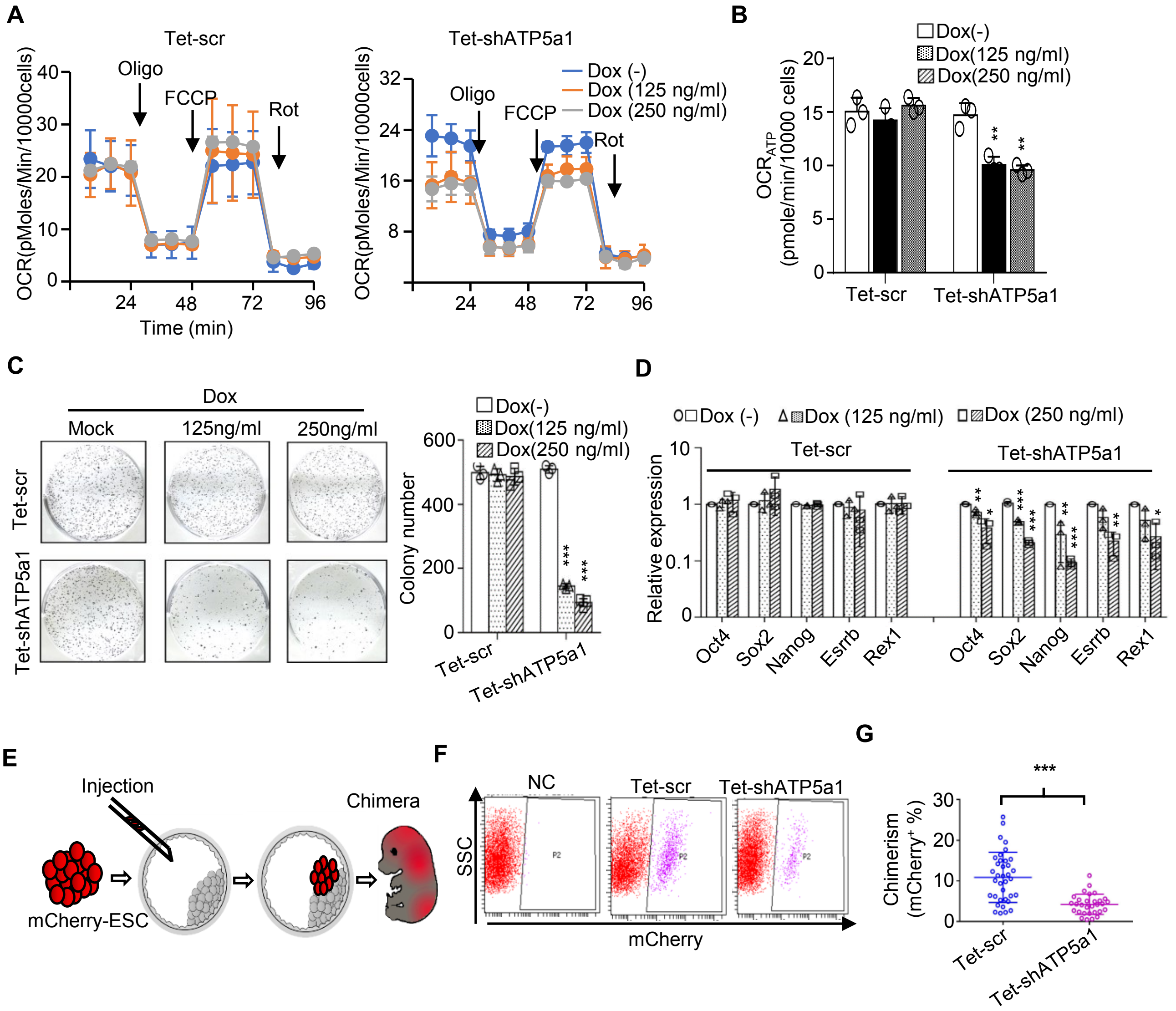


**Figure S6. Genetic Inhibition of Oxidative Phosphorylation Causes Loss of ESC Identity** (**A**) Oxygen consumption rate (OCR) measurements on stable ESC lines expressing either scramble shRNA or shRNA targeting ATP5a1 under Dox control. (**B**) The ATP production-related OCR of Tet-shATP5a1 ESC decreases upon Dox treatment. Results are shown as mean ± SD of triplicates from one representative of three independent experiments. $^{**}P < 0.01$; Student's *t*-test. (**C**) Knockdown of ATP5a1 inhibits ESC self-renewal. Left, representative photos of ESC colonies stained with alkaline phosphatase. Right, statistical analysis of alkaline phosphatase-positive colonies. Results are shown as mean ± SD from 3 independent experiments. $^{***}P < 0.001$; Student's *t*-test. (**D**) ATP5a1 knockdown decreases pluripotency gene expression. Results are shown as mean ± SD of 3 independent experiments. $^{*}P < 0.05$; $^{**}P < 0.01$; $^{***}P < 0.001$; Student's *t*-test. (**E**) Diagram of the chimeric mouse formation assay. (**F**) Knockdown of ATP5a1 decreases the ESC chimera rate. The mCherry-positive cells detected by FACS indicate the number of cells in each chimeric embryo that were derived from the originally injected cells. (**G**) Summary of data from chimeric embryos. Each dot represents the percentage of $mCherry^{+}$ cells in an individual chimeric embryo. Scr, n=36; ATP5a1, n=30. $^{***}P < 0.001$; Student's *t*-test.

**Figure S7. O-GlcNAcylation of Pluripotency Factors is Essential for ESC Identity**

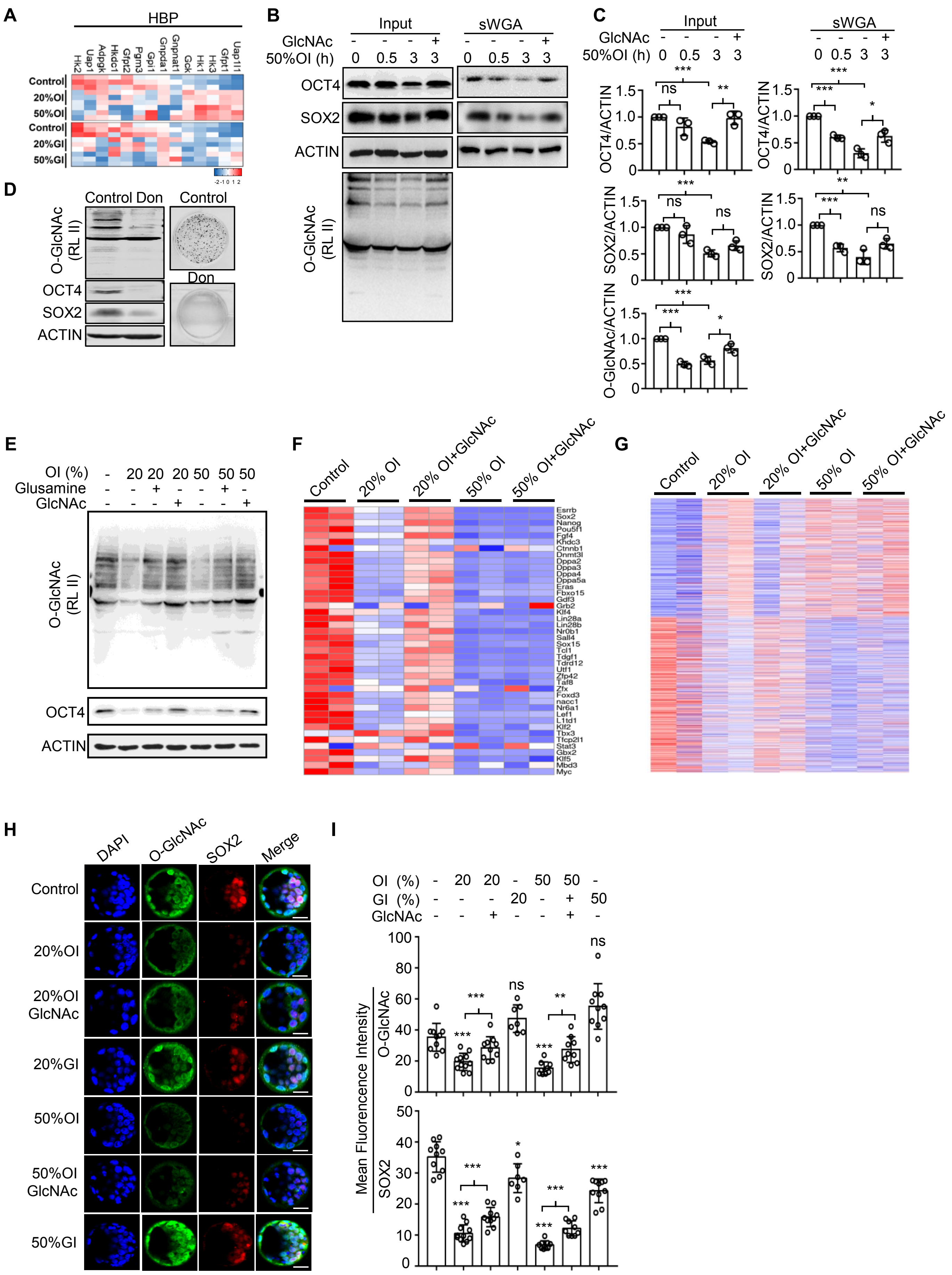

**Figure S7. O-GlcNAcylation of Pluripotency Factors is Essential for ESC Identity** (**A**) Heatmap of expression of genes involved in the HBP pathway. (**B**) 50%OI treatment causes decreased O-GlcNAcylation and expression of SOX2 and OCT4 in naïve-state ESCs. (**C**) Statistical analysis of the western blot results shown in **B**. n=3; *P < 0.05; **P < 0.01; ***P < 0.001; ns, not significant; Student's *t*-test. (**D**) Inhibition of the HBP pathway leads to breakdown of ESC identity. Left, HBP pathway inhibition by Don causes decreased expression of OCT4 and SOX2; right, representative images of ESC colonies stained with alkaline phosphatase. Don, 6-diazo-5-oxo-L-norleucine. (E) Both GlcNAc and glucosamine rescue the decreased O-GlcNAcylation and expression of OCT4 caused by OXPHOS inhibition. (**F**) Compromised expression of pluripotency genes led by OXPHOS inhibition is partially rescued by GlcNAc. Values displayed correspond to the expression level in the indicated sample scaled by the mean expression of each gene across samples. (**G**) OXPHOS inhibition causes altered expression of genes in the whole protein-coding transcriptome, and this effect is partially rescued by GlcNAc. Values displayed correspond to the expression level in the indicated sample scaled by the mean expression of each gene across samples. (**H**) Inhibition of OXPHOS rather than glycolysis reduces O-GlcNAcylation and expression of SOX2 in the inner cell mass of *ex vivo*-cultured blastocysts. These effects are ameliorated by adding GlcNAc. Confocal immunofluorescence microscopy was carried out using antibodies specific to O-GlcNAc (green) and SOX2 (red). The nuclei were stained with DAPI. Bars, 25 μm. (**I**) Statistical analysis of the immunofluorescence microscopy results shown in **H**. Control, n=10; 20%OI, n=11; 20%OI+GlcNAC, n=10; 20%GI, n=7; 50%OI, n=9; 50%OI+GlcNAC, n=9; 50%GI, n=10; *P < 0.05; **P < 0.01; ***P < 0.001; ns, not significant; Student's *t*-test.

**Figure S8. OXPHOS Inhibition Induces ESCs to A State Which is Different to Diapause- or Primed- State**

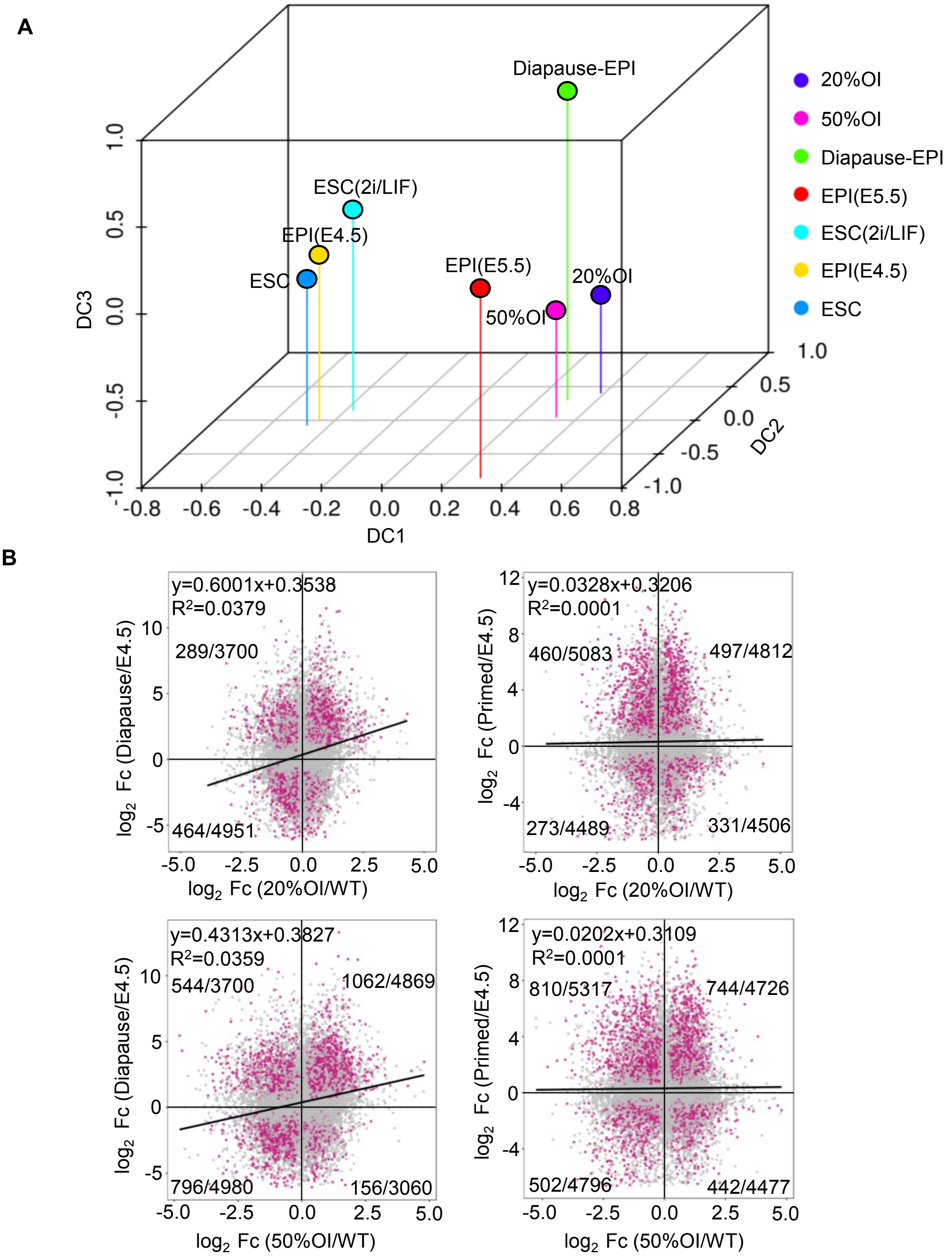


**Figure S8. OXPHOS Inhibition Induces ESCs into a State Which is Different to the Diapause- or Primed- State** (**A**) Diffusion map of ESCs, 20%OI- or 50%OI- treated ESCs and ESCs(2i/LIF), EPIs(E4.5), diapause EPIs or EPIs(E5.5) shows that OXPHOS inhibition induces ESCs into a unique state, which is closer to the diapause- state than to the primed- state (EPIs, epiblasts). (**B**) Scatterplots of gene expression levels that are significantly changed in the comparison of 20%OI/WT and Diapause/E4.5 (top left), 20%OI/WT and Primed(E5.5)/E4.5 (top right), 50%OI/WT and Diapause/E4.5 (bottom left) or 50%OI/WT and Primed(E5.5)/E4.5 (bottom right). The x axis shows the $\log_2$ fold change ($\log_2$ Fc) between 20%OI-(or 50%OI-) ESCs and control ESCs; the y axis shows the $\log_2$ Fc between diapause epiblasts (or E5.5 primed epiblasts) and E4.5 epiblasts. The dots represent the set of all genes. Differentially expressed genes ($|FC|\geq1.5$ and $p\leq0.05$) are shown as red dots. The numbers of DEGs (red) and total genes (gray) are labeled in every corresponding quadrant. The linear equation is fitted by all the dots in the four quadrants, and the slope of this linear fitting equation partially reflects the correlation of the x and y axes. $R^2$ values display correlation coefficients. Data for ESCs(2i/LIF), EPIs(E4.5), diapause EPIs and EPIs(E5.5) are from Boroviak et al. 2015.

**Figure S9. Network Integration of Metabolome and Transcriptome Data Reveals that Inhibition of OXPHOS Leads to Defective UDP-GlcNAc Biosynthesis**

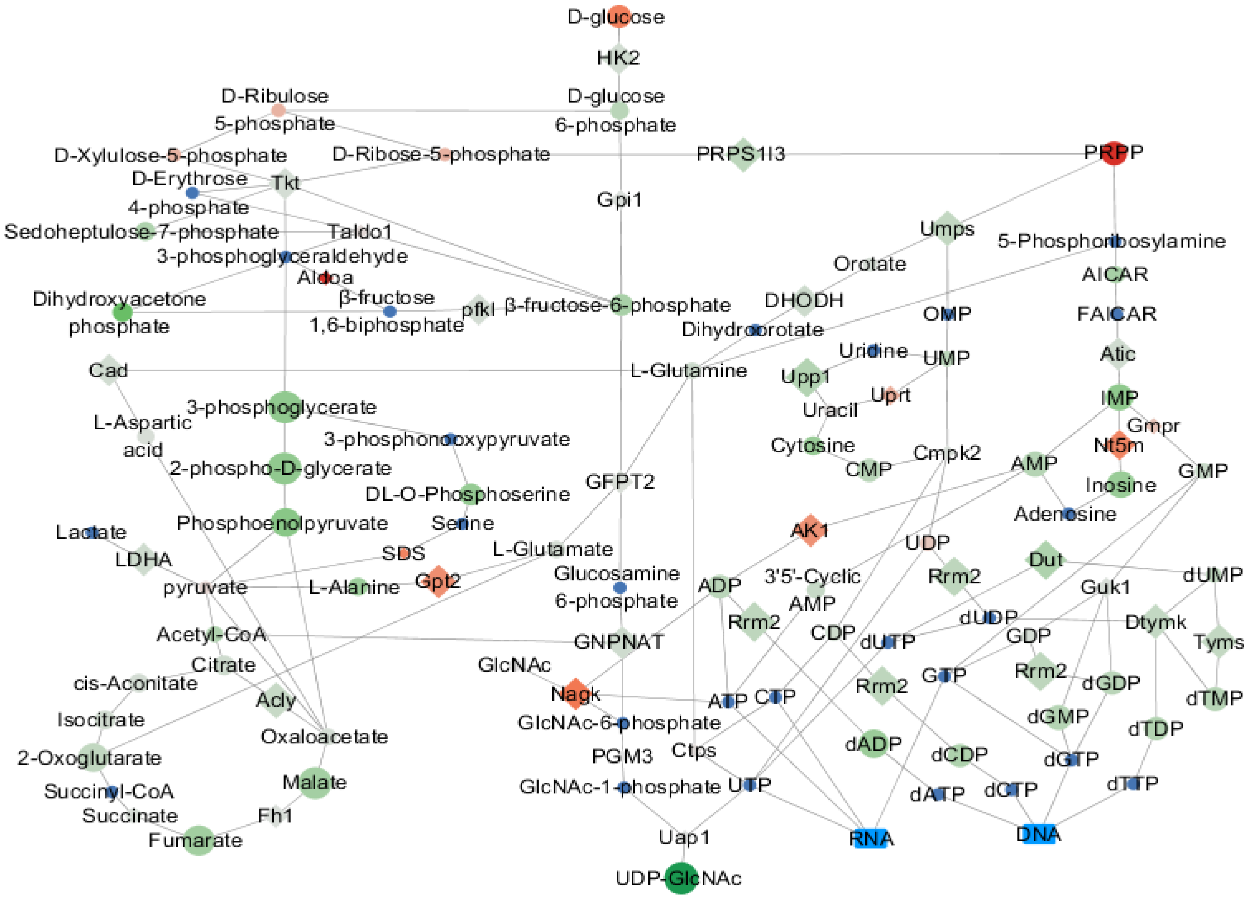


**Figure S9. Network Integration of Metabolome and Transcriptome Data Reveals that Inhibition of OXPHOS Leads to Defective UDP-GlcNAc Biosynthesis** A global combined network that connects metabolite nodes to reaction nodes was constructed based on the detected metabolites and the corresponding enzymes. The node of each reaction is connected to its respective substrates and products (solid circles). Enzymes that catalyze representative reactions are shown (diamonds). The relative levels of metabolites and enzymes in 20%OI ESCs compared to control ESCs is indicated by different colors (red, upregulated; green, downregulated; blue, not measured), and the significance is indicated by dot size (the bigger the more significant).

**Figure S10. UDP-GlcNAc Links Oxidative Phosphorylation to Pluripotency**

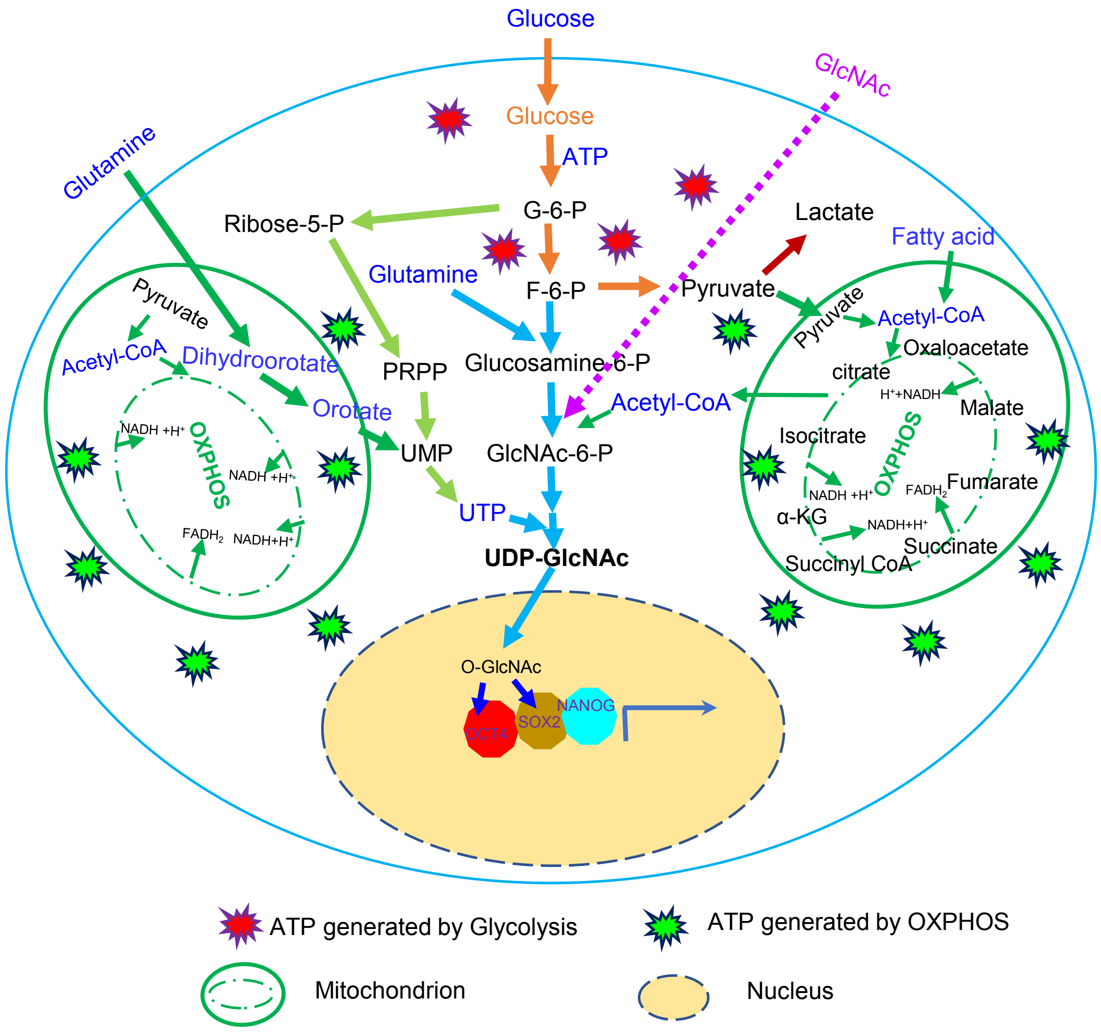


**Figure S10. UDP-GlcNAc Links Oxidative Phosphorylation to Pluripotency** ESC mitochondria have a high ATP generation capacity. Mitochondrial respiration produces the majority of cellular ATP in ESC, and couples with the hexosamine biosynthesis pathway to generate UDP-GlcNAc for O-GlcNAcylation of pluripotency factors like OCT4 and SOX2, thereby safeguarding ESC identity. Moderate inhibition of mitochondrial respiration leads to incomplete catabolism of glucose together with abnormal metabolism of glutamine, nucleotides, and acetyl-CoA, resulting in decreased UDP-GlcNAc generation and compromised pluripotency. This effect can be ameliorated by direct supplementation of GlcNAc.
