## Supplementary material for "Oxidative phosphorylation safeguards pluripotency via UDP-N-acetylglucosamine": Table S3 Key resources table

| REAGENT or RESOURCE | SOURCE | IDENTIFIER |
|---|---|---|
| Antibodies | | |
| Rabbit polyclonal OCT4 antibody | Abcam | Cat#ab19857; RRID:AB_445175 |
| Mouse monoclonal OCT4 antibody | Santa cruz | Cat#sc-5279; RRID:AB_628051 |
| Rabbit polyclonal SOX2 antibody | Millipore | Cat#AB5603; RRID:AB_2286686 |
| Rabbit polyclonal SOX2 antibody | Abcam | Cat#ab97959; RRID:AB_2341193 |
| Rabbit polyclonal Nanog antibody | Abcam | Cat#ab80892; RRID:AB_2150114 |
| Mouse monoclonal ATP5A antibody | Abcam | Cat# ab176569, RRID:AB_2801536 |
| Mouse monoclonal UQCRC2 antibody | Abcam | Cat# ab14745, RRID:AB_2213640 |
| Rabbit polyclonal TOM 40 antibody | Fine Test | Cat# FNab08860 |
| Rabbit polyclonal TIM 23 antibody | Fine Test | Cat# FNab08693 |
| Mouse monoclonal [RL2] to O-Linked N-Acetylglucosamine antibody | Abcam | Cat#ab2739; RRID:AB_303264 |
| Rabbit monoclonal [EPR12713] to OGT / O-Linked N-Acetylglucosamine Transferase | Abcam | Cat#ab177941 |
| Mouse Monoclonal Anti-β-Actin antibody | Sigma | Cat#A5441; RRID:AB_476744 |
| anti-mouse IgG (H+L), F(ab') 2 Fragment (Alexa Fluor 488 Conjugate) | Cell Signaling Technology | Cat#4408; RRID:AB_10694704 |
| anti-rabbit IgG (H+L), F(ab') 2 Fragment (Alexa Fluor 647 Conjugate) | Cell Signaling Technology | Cat#4414; RRID:AB_10693544 |
| HRP-conjugated goat anti-rabbit IgG | Beyotime | Cat#A0208 |
| HRP-conjugated goat anti-mouse IgG | Beyotime | Cat#A0216 |
| Bacterial and Virus Strains | | |
| Tet-pLKO-puro lentiviral vector | Wiederschain et al., Cell Cycle. 2009 Feb 1. 8(3):498-504 | Addgene: 21915; RRID:Addgene_21915 |
| pSPAX2 | Didier Trono | Addgene: 12260; RRID:Addgene_12260 |
| pMD2G | Didier Trono | Addgene: 12259; RRID:Addgene_12259 |
| Chemicals, Peptides, and Recombinant Proteins | | |
| Oligomycin | Sigma | Cat#O4876 |
| FCCP | Sigma | Cat#C2920 |
| Rotenone | Sigma | Cat#R8875 |
| Antimycin A | Abcam | Cat#ab141904 |
| 2-DG | Sigma | Cat#D8375 |
| Digitonin | Sigma | Cat#D141 |
| Pyruvate | Sigma | Cat#107360 |
| Malic acid | Sigma | Cat#02288 |

| | | |
|---|---|---|
| Adenosine 5′-diphosphate sodium salt | Sigma | Cat#A2754 |
| 6-Diazo-5-oxo-L-norleucine | Selleckchem | Cat#S8620 |
| TMRE | Thermos Fisher | Cat#T669 |
| Deep Red FM | Invitrogen | Cat#M22426 |
| Glucosamine hydrochloride | Sigma | Cat#G1514 |
| N-Acetyl-D-glucosamine | Sigma | Cat#A3286 |
| Puromycin | Gibco | Cat#A1113802 |
| Doxycycline | Sigma | Cat#D9891 |
| DiO | Yeasen | Cat#40725ES10 |
| BSA(Fatty Acid & IgG Free) | Beyotime | Cat#ST025 |
| $MgCl_2$ | Sigma | Cat#M2393 |
| $KH_2PO_4$ | Sigma | Cat#P5655 |
| HEPES | Sigma | Cat#H3375 |
| EGTA | Sigma | Cat#E3889 |
| Sucrose | Sigma | Cat#S7903 |
| D-Mannitol | Sigma | Cat#M4125 |
| Agarose Wheat Germ Agglutinin (WGA), Succinylated | Vector Laboratories | Cat#AL-1023S |
| Critical Commercial Assays | | |
| Alkaline Phosphatase Assay Kit | Beyotime | Cat#P0321 |
| Mitochondria Isolation Kit for Cultured Cells | Thermos Fisher | Cat#89874 |
| Luminescent ATP Detection Kit | Promega | Cat#ab113849 |
| RNeasy Mini Kit | Qiagen | Cat#74134 |
| GoTaq® qPCR Master Mix | Promega | Cat#PRA6101 |
| SuperScript™ III First-Strand Synthesis System | Invitrogen | Cat#18080051 |
| Deposited Data | | |
| Raw and analyzed data | This paper | GEO: GSE140712 |
| Experimental Models: Cell Lines | | |
| Mouse: B6 ESC | This paper | N/A |
| Mouse: 129 ESC | This paper | N/A |
| Mouse: mito-red ESC | This paper | N/A |
| Mouse: GFP ESC | This paper | N/A |
| Mouse: Tet-shATP5a1 | This paper | N/A |
| Mouse: Tet-scr | This paper | N/A |
| Mouse: B6 MEF | This paper | N/A |
| Mouse: 129MEF | This paper | N/A |
| Experimental Models: Organisms/Strains | | |
| Mouse: C57BL/6J | Jackson Laboratory | IMSR Cat# JAX: 000664; RRID:IMSR_JAX:000664 |
| Mouse: 129X1/SvJ | Jackson Laboratory | IMSR Cat# JAX: 000691; RRID:IMSR_JAX: 000691 |
| Mouse: B6D2-Tg(CAG/Su9-DsRed2,Acr3- | Riken | IMSR Cat# RBRC03743; |

| | | |
|---|---|---|
| EGFP)RBGS002Osb | | RRID:IMSR_RBRC03743 |
| Mouse: B6.Cg-Tg(CAG-GFP/LC3)53Nmz/NmzRbrc | Riken | IMSR Cat# RBRC00806; RRID:IMSR_RBRC00806 |
| Mouse: Nude mice | Charles River | NU/NU |
| Mouse: CF1 mice | SLACCAS | CF1 |
| Oligonucleotides | | |
| shRNA targeting ATP5a1 : 5'-TGCAGCCAAGATGAACGATTCCTTCAA GAGAGGAATCGTTCATCTTGGCTGCTTT TTTC-3’ | This paper | N/A |
| scramble shRNA : 5'-TGCTTACGCTGAGTACTTCGAATTCAA GAGA TTCGAAGTACTCAGCGTAAGCTTTTTTC-3’) | This paper | N/A |
| Primers for real-time PCR, see Materials and Methods | This paper | N/A |
| Recombinant DNA | | |
| p-Tet-scr | This paper | N/A |
| p-Tet-shATP5a1 | This paper | N/A |
| pCDH-CAG- TetR-ires- mcherry | This paper | N/A |
| pCDH-CAG-mcherry | This paper | N/A |
| Software and Algorithms | | |
| Prism 8 | GraphPad Software | https://www.graphpad.com/scientific-software/prism/ |
| Cytoscape | ShannonP et al., 2013 | https://cytoscape.org/ |
| Imaris 9.2.1 | OXFORD Instruments | https://imaris.oxinst.com |
| Rstudio | Allaire et al., 2012 | https://rstudio.com/ |
| ImageGP | EHBIO gene technology | http://www.ehbio.com/ImageGP/ |
| ImageJ | National Institutes of Health | https://imagej.nih.gov/ij/ |
| Other | | |
