## Supplementary material for "Oxidative phosphorylation safeguards pluripotency via UDP-N-acetylglucosamine": pdf Video legends

Video S1: Representative 3D-SIM image of a mouse ESC. Red, mitochondrion; Green, cell membrane

Video S2: Representative 3D-SIM image of a mouse EpiLC. Red, mitochondrion; Green, cell membrane

Video S3: Representative 3D-SIM image of a mouse NSC. Red, mitochondrion; Green, cell membrane

Video S4: Representative 3D-SIM image of a mouse MEF. Red, mitochondrion; Green, cell membrane

Video S5: Representative 3D-SIM image of a mouse HL-1 cell. Red, mitochondrion; Green, cell membrane
